# The ZIP8 A391T Crohn’s disease-linked risk variant induces colonic metal ion dyshomeostasis, microbiome compositional shifts, and inflammation

**DOI:** 10.1101/2024.08.19.608695

**Authors:** Julianne C. Yang, Matthew Zhao, Diana Chernikova, Nerea Arias-Jayo, Yi Zhou, Jamilla Situ, Arjun Gutta, Candace Chang, Fengting Liang, Venu Lagishetty, Jonathan P. Jacobs

## Abstract

The pathogenesis of Crohn’s disease involves genetic and environmental factors, with the gut microbiome playing a crucial role. The Crohn’s disease-associated variant rs13107325 in the SLC39A8 gene results in an A391T substitution in the ZIP8 metal ion transporter and has previously been linked to alterations in the colonic microbiome in variant carriers. We hypothesized that the A391T substitution alters metal ion homeostasis in the colonic mucosal-luminal interface, thereby inducing dysbiosis which may promote intestinal inflammation. To evaluate this hypothesis, we generated a SLC39A8 A393T mouse model (matching human A391T). Consistent with an effect of the variant on ZIP8 function, homozygous A393T mice exhibited increased cobalt in the colonic mucosa, but reduced iron, zinc, manganese, cobalt, copper, and cadmium in the colonic lumen. We performed 16S rRNA gene sequencing of colon samples and histological scoring of colon tissue collected from the SLC39A8 A393T mouse model at different ages. We identified variant-linked effects on microbiome beta diversity in 2-month, 3-4 month, and 12-month old mice and spontaneous intestinal inflammation in 10-month but not 5-month old mice. Predicted pathway analysis of the microbiome samples revealed differential enrichment of iron-, zinc- and cobalt-dependent pathways in A393T mice compared to wild-type controls. These results suggest that the variant in SLC39A8 primarily restricts metal availability to the microbiota, resulting in compositions that can adapt to the environment, and that A393T-linked dysbiosis occurs prior to the onset of inflammation. This study paves the way for future studies investigating risk variants as microbiome-disease modifiers.

**NEW & NOTEWORTHY:** Mice recapitulating the human A391T Crohn’s disease-linked risk variant in the SLC39A8-encoded ZIP8 transporter exhibit reduced luminal trace metals and an altered microbiome composition. The microbiome compositional shift occurs prior to the onset of spontaneous colonic inflammation. This study supports an emerging paradigm by which genetic risk for inflammatory bowel disease may confer disease susceptibility through modulation of the gut microbiome.

## INTRODUCTION

Crohn’s disease (CD) is an inflammatory bowel disease (IBD) characterized by inflammation along any part of the gastrointestinal tract, though it most commonly occurs in the ileocecal region. Both genetic and environmental influences are thought to contribute to the pathogenesis of Crohn’s disease. CD-associated genetic risk factors have been reported through genome-wide association studies (GWAS); among these are variants in genes which regulate the immune response such as *NOD2*, *CARD9*, and *REL*, and variants in genes which mediate tolerogenic responses to commensals, such as *IL10*, *IL27*, and *CREM*^1^. Lifestyle factors, such as smoking and diets high in saturated fat, have also been demonstrated to increase the risk of CD ^2^. This has led to increasing attention to the possible involvement of the CD-associated gut microbiome as a vector of both genetic and environmental insults in mediating colitis. IBD risk variants, including NOD2, have been associated with altered gut microbiota composition in humans ^3^. Therefore, interrogating whether CD risk variants can be disease-driving “microbiome quantitative trait loci” (mb-QTL) may be important to understanding the relationship between variant carrier status and heterogeneity in susceptibility to IBD ^4^.

Of the 200+ identified IBD risk variants from GWAS, the majority are non-coding, adding an additional layer of difficulty for mechanistic research in preclinical models ^5^. We and others have recently reported that the coding single nucleotide polymorphism (SNP) rs13107325 in the gene *SLC39A8* resulting in the Ala391Thr substitution in the ZIP8 metal ion transporter is associated with Crohn’s disease ^6^ ^7^. We additionally reported that Thr allele carriage was associated with altered microbiome composition in mucosal lavage samples from both CD patients and their healthy controls, and microbes depleted in healthy control carriers overlapped with those depleted in CD patient carriers ^6^. Since rs13107325 is also associated with other conditions spanning several organ systems, we hypothesized that this SNP may induce a diverse range of outcomes through its effects on the intestinal gut microbiome ^8^.

Two studies have since reported that A393T mice recapitulating the A391T human variant in *SLC39A8* exhibit increased intestinal inflammation in the dextran sulfate sodium (DSS) model of colitis. Both studies raised manganese tissue dyshomeostasis as a potential mechanism by which variant carrier status influences susceptibility to DSS-mediated colitis; however, conclusions on whether A393T carriers exhibit differential Mn abundance in the colon compared to their wild-type (WT) controls differed ^7^ ^9^. These studies also did not address the impact of the variant on the colonic microbiome. Therefore, in this study, we sought to clarify the role of the *SLC39A8* variant on ZIP8 transport function at the colonic luminal-mucosal interface, on the colonic microbiome, and on spontaneous and chemically-induced inflammation, and identify possible means by which these factors interact.

## MATERIALS AND METHODS

### Construction of the SLC39A8 A393T mouse model

SLC39A8 A393T knock-in mice were created through fertilized oocyte injection of Cas9 DNA endonuclease, two guideRNA molecules [CTAGCTTTCGGCATTTTGGT] and [AGCGAGTGCAAATATAATAT] targeting the region encoding Ala393 in SLC39A8, and a single-stranded oligodeoxynucleotide donor template containing three point mutations corresponding to T393 in addition to silent mutations preventing further Cas9 mediated mutation (**Figure S1**). The injected, fertilized oocyte was then transferred into a pseudopregnant female. Sanger sequencing was used to validate successful gene knockin of the offspring, which were subsequently backcrossed for three generations onto the C57Bl/6 background.

### Animals

Mice were given food (LabDiet 5053; irradiated) and water *ad libitum* and housed in static cages with autoclaved bedding under a 12:12 light:dark cycle. Colon samples were harvested from one cohort of 5-month old mice for inductively-coupled mass spectrometry (Figure 2) and for histological scoring (Figure 6A). For microbiomics sequencing, colonic luminal and mucosal samples were collected from one cohort of 2-month old mice and one cohort of 12-month old mice using a previously described protocol^15^. Fecal pellets from a separate cohort of 3-4 month old mice were also collected and utilized for microbiomics sequencing. Mucus layer thickness was assessed in another cohort of 5-6 month old mice. Two cohorts of mice aged 9-12 weeks were used for 2,4,6-Trinitrobenzene sulfonic acid (TNBS) or dextran sodium sulfate (DSS)-mediated chemical induction of colitis.

### Mucus layer assessment

Mucus layer staining was performed using alcian-blue periodic acid-Schiff staining of paraffin-embedded, Carnoy’s fixed colon tissue. Staining was performed using the Richard-Allan Scientific Alcian Blue pH 2.5 Periodic Acid-Schiff Stain Kit according to the manufacturer’s protocol. Stained sections were assessed under light microscopy, and four images per animal were captured for mucus layer measurement. Inner mucus layer thickness was assessed using the Java-based image-processing program ImageJ. After calibration with a hemocytometer grid, three measurements were taken per image, for a total of twelve measurements averaged per animal.

### Histological scoring

Hematoxylin and eosin stained sections of colon were scored for the severity of intestinal inflammation as described by Erben et al ^16^. Scoring was performed by a single reviewer in a blinded manner. Histological scores ranged from 0 to 5, where 0=none, 1=minimal, 2=mild, 3=moderate, and 4/5=marked. Scores were based on the observed level of inflammatory cell infiltration, epithelial changes, and mucosal architecture disruption. Inflammatory cell infiltration limited to the mucosa corresponded to an overall score of 1, any extent of submucosal infiltration corresponded to an overall score of 2-4, and transmural inflammation corresponded to an overall score of 5. Epithelial changes were assessed based on the level of epithelial hyperplasia, as well as the extent of goblet cell loss and presence of crypt abscesses. Mucosal architecture was assessed, in which the presence of crypt irregularity or loss, with or without ulcerations, corresponded to an overall score of 4-5.

### Dextran sodium sulfate (DSS) chemical induction model of colitis

Mice were administered 2% DSS (MP Biomedicals cat # MFCD00081551) in drinking water for 7 days and switched to normal drinking water for 3 days. Body weights of the mice were obtained on the day prior to DSS administration and daily up to day 10 to calculate body weight as a percentage of baseline weight as the experiment’s primary outcome. Following euthanization on day 10, colons were harvested and measured, since colon length shortening is an additional indicator of inflammation in the DSS model. ^17^

### Trace element assessment via inductively-coupled plasma mass spectrometry (ICP-MS)

To reduce the effects of recent intake on the intestinal ion concentrations, food and water was withheld from the mice for 4 hours prior to euthanasia. Subsequently, the colon and the cecum were harvested. Luminal contents were collected via gentle squeezing of the intestines. Intestines were then cut lengthwise. One longitudinal half was cleaned in deionized water for ten seconds, which later became the “bulk colon tissue” samples for ICP-MS. The other longitudinal half was cleaned in phosphate buffered saline (PBS), then shaken vigorously in 15 mL of DMEM/1 mM dithiothreitol for 20 minutes at 37 °C. After shaking, an additional 15 mL of DMEM was added to the 50 mL conicals containing the samples, then the samples were vortexed and poured through a 70 µM filter. The samples were then centrifuged at 2000 xg for 20 minutes, decanted, resuspended in 15 mL DMEM, centrifuged at 2000 xg for 20 minutes,and then decanted. The pellets were then resuspended in 15 mL PBS, centrifuged at 700 xg for 8 minutes at 4 °C, then decanted, resuspended in 2 mL PBS, transferred to an Eppendorf tube, and then centrifuged for the last time at 700 xg for 8 minutes. Remaining PBS was removed with a pipette, taking care not to disturb the pellet. All centrifugation was performed at 4 °C. The colonic luminal, cleaned colon tissue, and mucosal epithelial cell pellets were then dried at 60 °C for 16 hours to obtain the dry weight. ICP-MS (NexION 2000, PerkinElmer) analysis was performed to detect multiple elements (Fe, Co, Cu, Zn, Cd, Mn) in the samples. All samples were used as received without further purification or modification. Each sample was transferred to a clean Teflon vessel for acid digestion. Digestion was carried out with concentrated HNO_3_ (65-70%, Trace Metal Grade, Fisher Scientific) with a supplement of H_2_O_2_ (30%, Certified ACS, Fisher Scientific) at 190 °C for 20 min in a microwave digestion system (Titan MPS, PerkinElmer). Once the sample was cooled to room temperature, it was subsequently diluted to make a final volume of 50 mL by adding filtered deionized water for analysis. The calibration curve was established using a standard solution while the dwell time was 50 ms with thirty sweeps and three replicates with background correction.

### 2,4,6-Trinitrobenzene sulfonic acid (TNBS) chemical induction model of colitis

This protocol was adapted from Wirtz et al ^18^. Briefly, mice were presensitized one week prior to intrarectal administration with 2.5% TNBS. For presensitization, a 1% TNBS solution was applied to a shaved 1.5 x 1.5 cm spot on the backs of the mice. After 7 days, 2.5% TNBS in ethanol was administered intrarectally with a rubber-tipped needle into anesthetized mice. The mice were held by the tail, snout downwards, for an additional minute to ensure proper administration. Mice were weighed for 3 days afterwards, and the lengths of colons were measured following euthanasia.

### 16S rRNA v4 gene sequencing

DNA was extracted from the colonic luminal content of 2-month old mice and 1-year-old mice utilizing the Zymobiomics DNA Miniprep Kit (cat#D4300) according to the manufacturer’s protocol. For fecal pellet samples from 3-4 month old mice, DNA extractions were performed with the Qiagen DNeasy PowerSoil Pro Kit (cat#47014). Next, to amplify the v4 region of the 16S rRNA gene, polymerase chain reactions (PCR) were set up for each sample using the extracted DNA as the template and primers 515F and 806R as previously described ^19^. PCR products were then purified with the ZR-96 DNA Clean & Concentrator-5 (cat#D4024) kit and pooled together for paired-end sequencing with Illumina NovaSeq. Following quality filtering and merging of raw reads utilizing the DADA2 package in R, a count table of unique amplicon sequence variants (ASV) x samples was obtained for each cohort (2-month, 3-4 month, and 1-year old). Taxonomy assignment of the ASVs was achieved using the naive Bayes classifier trained on the Silva 138.1 database. For the 2-month and 1-year cohorts, the datasets were stratified by sample type (Luminal or Mucosal) prior to further analysis. The 2-month old mice were sequenced in two batches (encoded by the variable Sequencing_Run). The 2-month old and 1-year old cohorts included luminal and mucosal samples collected from the cecum, proximal colon, and distal colon (encoded by the variable Site), whereas the 3-4 month old samples were sequenced in one batch with one fecal sample per mouse. Calculation of square-root Jensen-Shannon dissimilarity matrices on the ASV count tables prevalence filtered to 15% followed by principal coordinates analysis for visualization was used to generate Figure 4. Repeated-measures PERMANOVA on the dissimilarity matrix was used to determine the effect of Genotype, after accounting for Sex (for the 3-4 month old mice), or after accounting for Site and Sex (for the 1-year old mice), or after accounting for Sequencing_Run, Site, and Sex (for the 2-3 month old cohort). For differential taxon association testing, ASV counts were first collapsed to the genus level. Next, the log-transformed, total sum scaling-normalized, 15% prevalence filtered genus-level counts were fitted to linear mixed-effects models (LMEM) implemented in MaAsLin2, with Sex and Genotype as fixed effects (3-4 month), Site, Sex, and Genotype as fixed effects and MouseID as a random effect (1-year), and Sequencing_Run, Site, Sex, and Genotype as fixed effects with MouseID as a random effect (2-month) ^20^. A genus was considered as significantly differentially abundant if, following multiple hypothesis correction, the q-value was less than 0.25. The PICRUSt2 algorithm was used to generate predicted pathway abundances based on the 16S data ^14^. Subsequently, the same LMEMs were fitted to the pathway abundance data in order to identify pathways differentially enriched in MUT compared to WT mice following multiple hypothesis correction (q<0.25). Circle plots grouping enriched pathways into higher-order hierarchies defined by MetaCyc were generated with package “circlize” in R. Analysis scripts and input files sufficient to replicate the figures (with the exception of Figure 1) are publicly available on https://github.com/jacobslabucla/slccolon/.

## RESULTS

### The SLC39A8 variant affects predicted ZIP8 structure and disrupts trace element homeostasis in the colon

Given that the SNP rs13107325 results in the substitution of the nonpolar Ala residue to the polar Thr residue, we reasoned that this could affect the ZIP8 protein structure and lead to functional effects on ion transport. Since the structure of ZIP8 has not yet been determined experimentally, we utilized the AlphaFold2 structural prediction algorithm to generate and overlay the predicted structures of human ZIP8 A391 and ZIP8 T391 ^10^. Though the distance between Ala and Thr at the 391 position was only 1.35Å as measured between alpha carbons, key glutamine and histidine residues which are purported to mediate metal ion binding in ZIP transporters were shifted by 3.49Å and 3.83Å in the structural overlay (**Figure 1**) ^11^.

**Figure 1.**
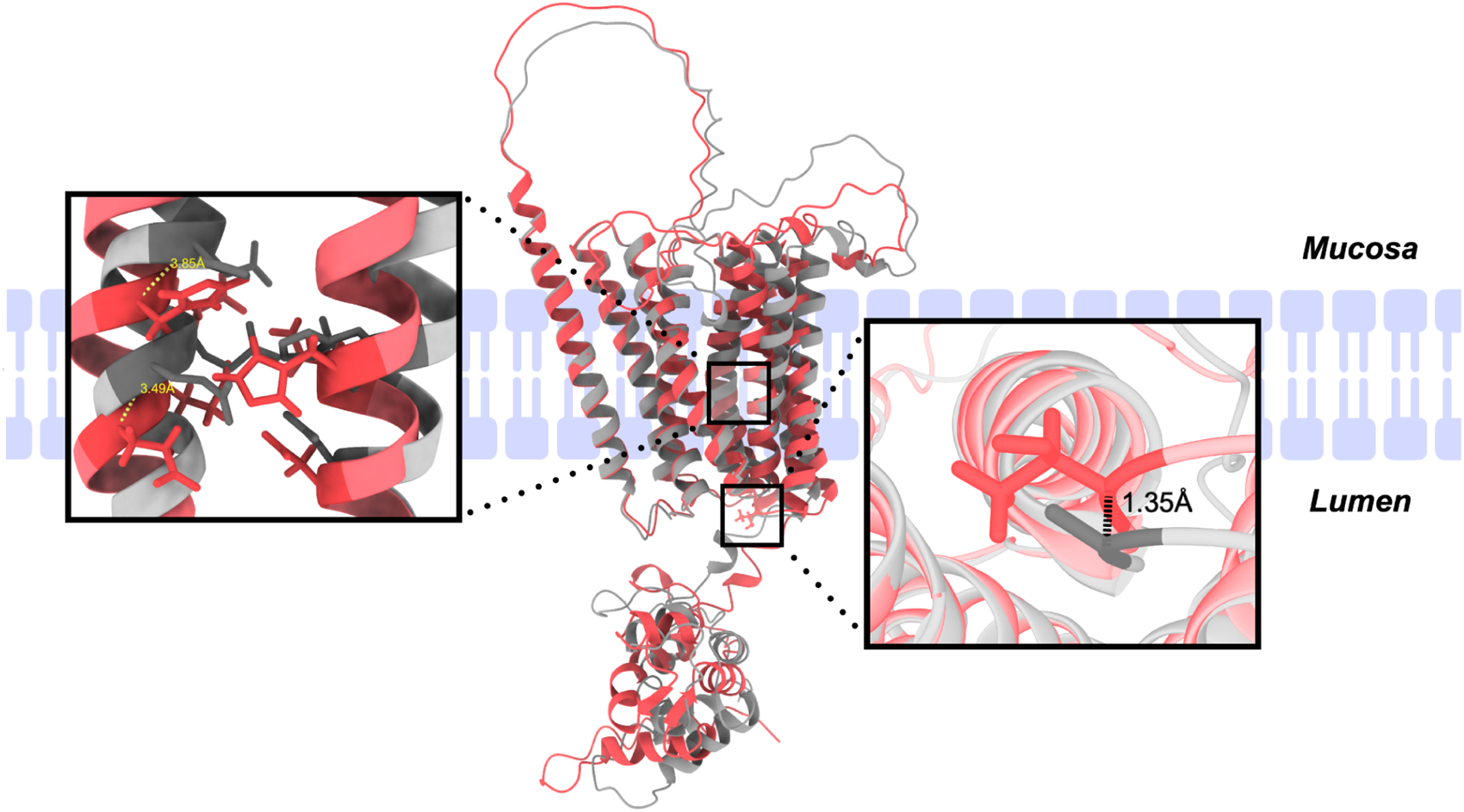
Overlay of ZIP8 A391 and ZIP8 T391 predicted structures. ZIP8 A391 (gray) and ZIP8 T391 (red) structures were generated with the AlphaFold2 structural prediction algorithm and overlaid in UCSF ChimeraX. The right box highlights the distance between alpha carbons in the residues at the 391 position. The left box highlights a EEXXHE motif on transmembrane domain V and HNXXD motif on transmembrane domain IV.

To evaluate whether the changes in ZIP8 structure were accompanied by changes in function, we generated ZIP8 A393T mice recapitulating the human variant by CRISPR-Cas mediated gene editing followed by three rounds of backcrossing. We then examined trace element abundance in colonic luminal, mucosal, and cleaned bulk tissue samples collected from WT mice and homozygous (MUT) A393T mice with inductively-coupled plasma mass spectrometry (ICP-MS). Trace elements evaluated included iron, cobalt, copper, zinc, cadmium, and manganese (**Figure 2**). Compared to WT mice, MUT mice exhibited significantly lower levels of the above elements in the colonic luminal samples, higher cobalt abundance in the mucosal epithelium, and higher cobalt, cadmium, and manganese in the cleaned bulk tissue (p<0.05 for all elements) (**Figure 2**). These results suggest that the functional consequences of ZIP8 A391T include increased uptake of these dietary elements by the host, and conversely, reduced bioavailability of these elements for the intestinal microbiota.

**Figure 2.**
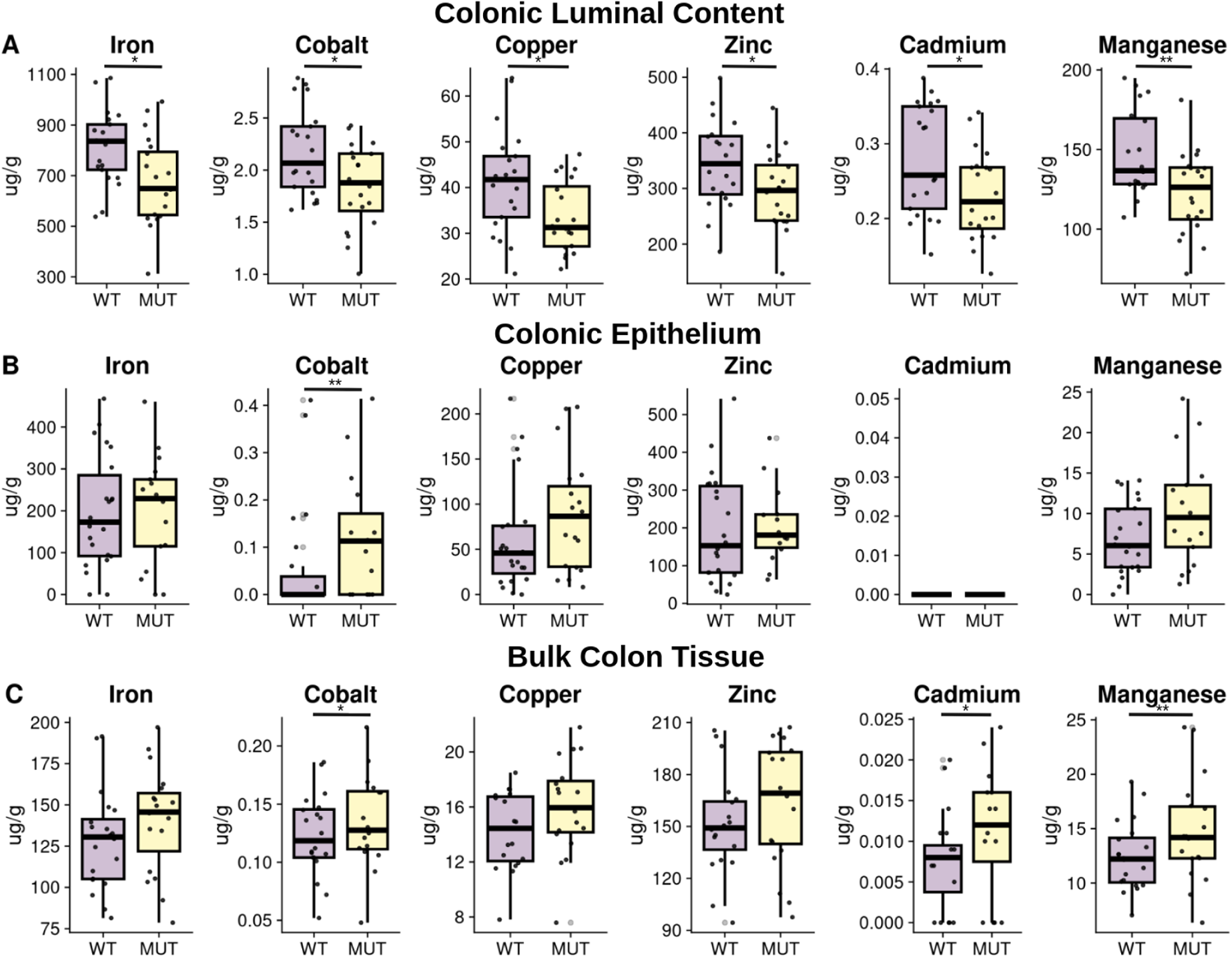
MUT mice exhibit altered luminal:mucosal trace element distributions compared to their WT counterparts. Harvested samples from 5-month old WT (n= 13 females (F) and n=12 males (M)) and MUT (n=12 F and 11 M) mice were utilized for trace element analysis via inductively-coupled plasma mass spectrometry (ICP-MS). Box plots show the distributions of trace element abundance by genotype in units of µg/g dry weight for colonic luminal content (**A**), colonic mucosa (epithelial cells) (**B**), and bulk colon tissue (**C**), with outliers removed according to the 1.5 InterQuartile Range criteria for visualization purposes only. Statistical analysis was performed on the whole dataset using Wilcoxon rank-sum tests comparing WT to MUT; *p<0.05; **p<0.01.

### The SLC39A8 variant induces microbiome compositional and predictive functional shifts that increase with age

Iron, copper, zinc and manganese are critical to the survival and fitness of many bacterial organisms, while cobalt and cadmium exposure have been reported to affect the gut microbiome ^12^ ^13^. Since we observed ∼25% reduction in all of these trace elements in the luminal samples of MUT compared to WT mice, we hypothesized that this would affect the gut microbiome of variant carriers. To thoroughly evaluate effects of the variant on the gut microbiome, we performed 16S rRNA gene sequencing of samples collected from three cohorts of mice: colonic luminal and mucosal-adherent microbiota of 2-month and 12-month old WT, heterozygous (HET), and MUT mice, along with the fecal microbiota of 3-4-month old WT, HET, and MUT mice. Due to the feces being representative of the contents in the distal end of the colon, we compared the 3-4 month cohort to the 2- and 12-month colonic luminal samples.

In the 2-, 3-4, and 12-month old luminal/fecal content, relative abundances of highly abundant bacteria were similar among the WT, HET, and MUT mice (**Figure 3A**, **3C**, and **3E**). However, visualization of microbiome beta-diversity through principal coordinates analysis (PCoA) of Jensen-Shannon distances illustrates MUT sample clustering along the PC1 axis (**Figure 3F**, **3H**, and **3J**). In all three luminal/fecal cohorts, PERMANOVA of the Jensen-Shannon distance matrix revealed that genotype (encoding WT, HET, or MUT) was significantly associated with microbiome composition, with R^2^ (effect size) values increasing from 3% in the 2-month cohort to 9% in the 12-month cohort (**Figure 3F**, **3H**, and **3J**).

Similarly, the 2- and 12-month old colonic mucosal samples did not show differences in highly abundant bacteria among the WT, HET, and MUT mice in the taxonomic summary plots, with the exception of reduced relative abundance in *Pseudomonas* of HET and MUT compared to WT 2-month old colonic mucosa and reduction in *Ligilactobacillus* in MUT compared to WT 12-month old colonic lumen and mucosa (**Figure 3B** and **3D**). While differences in mucosal colon community structure by genotype were only marginally significant for the 2 month old mice, the association of genotype with gut microbiome was statistically significant with an R^2^ of 6% in the 12 month old mice (**Figure 3I**). There were no differences in microbiome alpha diversity as assessed through the Shannon index (species richness and evenness) or by total number of amplicon sequence variants (ASVs) across genotypes for any of the datasets with the exception of increased ASVs in MUT compared to WT colonic mucosal samples (**Figure S2**). Overall, these findings support an effect of variant carrier status on microbiome beta-diversity which increases with age and is greater in the luminal compartment of the colon than in the mucosal-adherent fraction.

**Figure 3:**
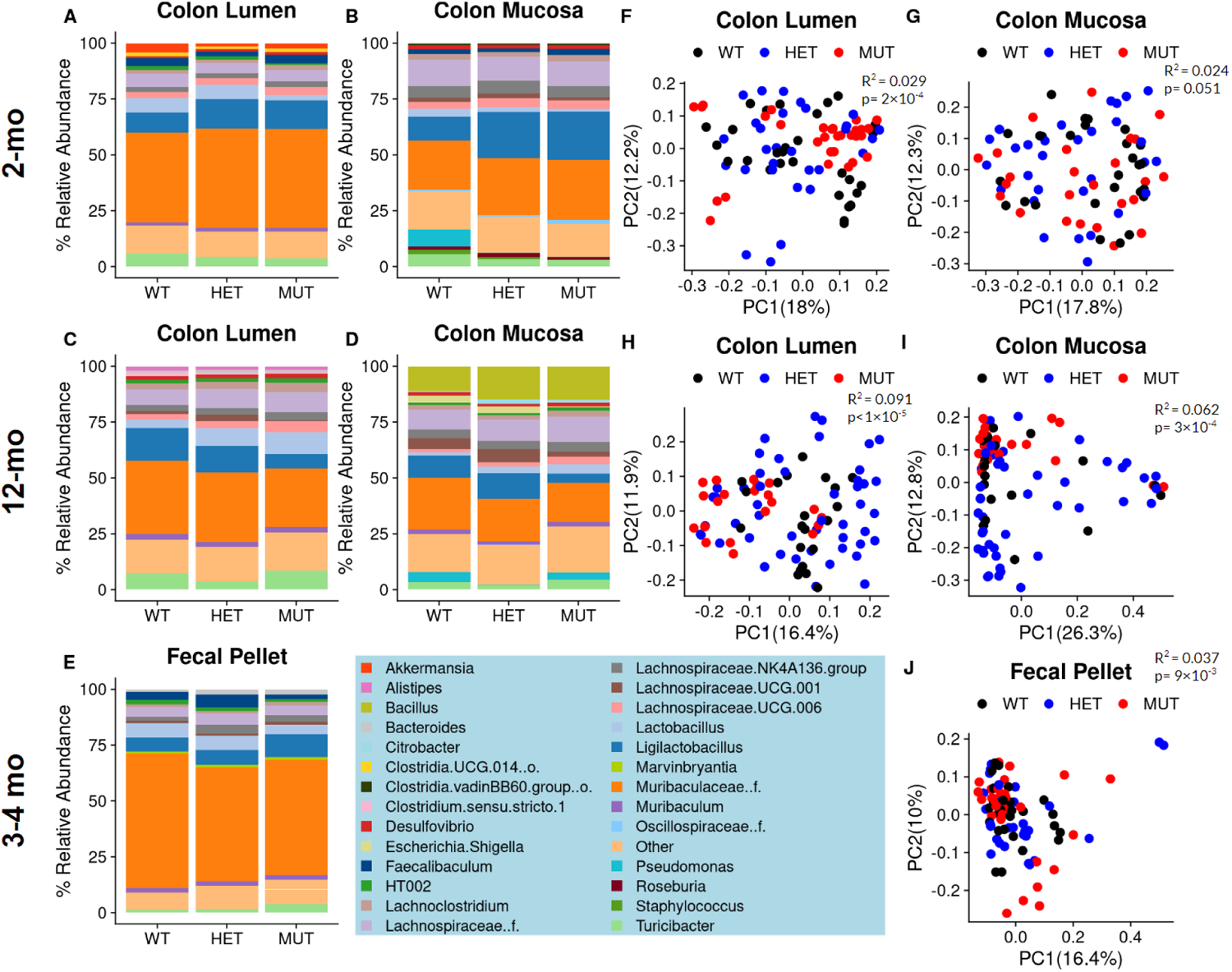
MUT mice exhibit a distinct microbiome composition in three differently aged cohorts. Taxonomic column summary plots illustrate the % relative abundance of genera which comprise at least 1% of the microbiome composition in the colonic lumen or adhered to the colonic mucosa of 2 month old WT, HET, and MUT mice (A, B), 12 month old mice (C, D), or the fecal pellets of 3-4 month old mice (E). The genera color legend for all barplots is shown at the bottom of the figure. Principal coordinates analysis plots on square-root Jensen-Shannon dissimilarities illustrate the differences in microbiome composition between WT, HET, and MUT mice, with each dot representing one sample. Plots are shown for the colonic lumen or colonic mucosa of 2-month old (F, G), 12-month old (H, I), or the feces of 3-4 month old (J) mice. R^2^ and p-values associated with the effect of Genotype are reported from PERMANOVA of sample dissimilarity matrices. The microbiome data represents n=10 WT (4 F 6 M), n=10 HET (4 F 6 M), and n=10 MUT (4 F 6 M) for 2 month-old mice, and n=27 WT (11 F 16 M), n=30 HET (10 F 20 M), and n=31 MUT (14 F 17 M) for 3-4 month-old mice.

Next, to assess whether changes in the abundance of specific genera accompany the MUT-linked differences in microbiome composition, we performed differential genera abundance testing comparing MUT to WT mice. While only 5-7 differentially abundant genera were found in the 2 and 3–4 month old mice, 18 and 29 differentially abundant genera were found in the 12 month old mice (**Figure 4**). Most differentially abundant genera had a relative abundance of <1%, with the exception of *Lactobacillus* and *Ligilactobacillus* in the 2- and 12-month old mice, respectively, and genera belonging to *Muribaculaceae* in the 3-4 month old mice (**Figure 4A**-**D**). Compared to their WT counterparts, *Lactobacillus* was significantly reduced in the 2 month old MUT mice in the colonic lumen, *Ligilactobacillus* was decreased in the the luminal and mucosal colons of the 12 month old MUT mice (**Figure 4B** and **4D**), and Muribaculaceae was significantly reduced in the 3-4 month old MUT mice (**Figure 4A-D**). Interestingly, *Staphylococcus*, *Rikenella*, *Akkermansia*, and *Candidatus_Arthomitus* were among 7 shared genera which were enriched in MUT compared to WT mice between both luminal and mucosal compartments of 12-month old mice, while no such overlap was observed in 2-month old mice (**Figure 4A**-**D**). Notably, across the three cohorts of mice, no genera were found to be differentially abundant in the same direction. These results suggest that variant-linked perturbations in microbiome community increase with age and may be due to broad shifts in multiple low-abundance bacterial taxa, rather than differential abundance of any particular bacteria.

**Figure 4:**
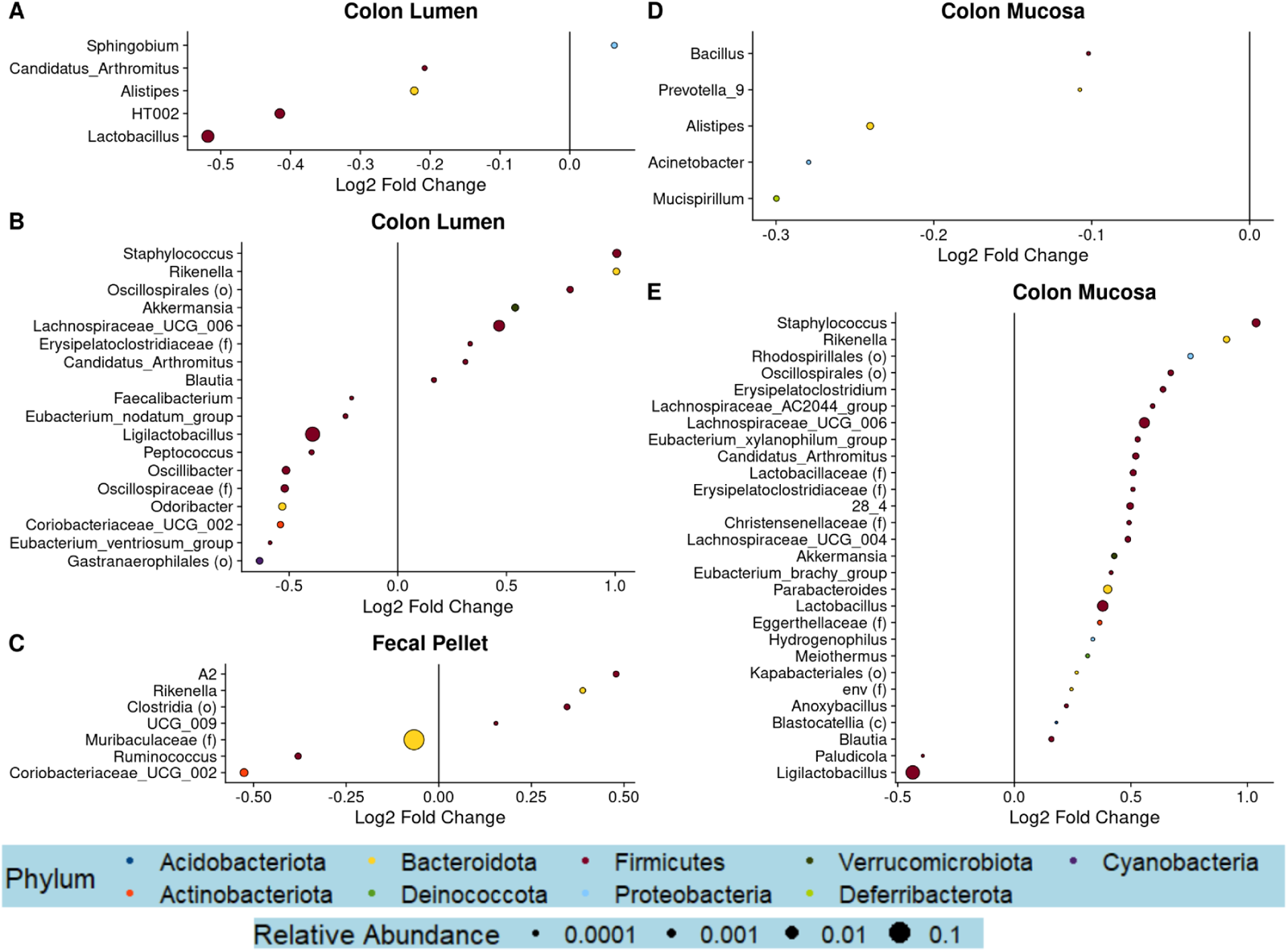
MUT mice exhibit altered abundance of several genera compared to WT mice. Differential abundance testing was performed at the genus level utilizing linear mixed-effects models of log-transformed, normalized count data. Dot plots visualizing significantly differentially abundant genera (q<0.25) between MUT and WT are shown for the colonic lumen or colonic mucosa datasets of the 2-month old mice (A, D), the 12-month old mice (B, E), or the fecal pellets of the 3-4 month old mice (C). The size of the dot corresponds to the relative abundance of the genera, while the color of the dot represents the phylum.

Since we did not identify genera which were consistently linked to variant carrier status across the three cohorts of mice, we investigated whether there were consistent patterns in microbial function that would better characterize the effects of the variant on the microbiome. To accomplish this, we utilized the PICRUSt2 pipeline to predict MetaCyc pathway abundances from the 16S rRNA gene sequencing count data ^14^. Predicted functional pathways which were identified as being significantly differentially enriched were present in all three cohorts. To identify patterns amongst the differentially enriched pathways, we grouped each individual pathway (bottom half of circular map) into higher-order classifications (top half of maps) (**Figure 5**). Interestingly, pathways classified as “cofactor biosynthesis” were observed in five out of six datasets, with the exception of the colonic lumen dataset in 2-month old mice (**Figure 5, Table S1**). In 2-month old mice, all 10 differentially abundant pathways classified as “cofactor biosynthesis” were depleted among mucosal-adherent bacteria of MUT mice compared to WT mice (**Figure 5B**, **Table S2**). In the 3-4 month old cohort, pathways classified as “cofactor biosynthesis” were both depleted and enriched, while in 12-month old mice, all 8 and 5 “cofactor biosynthesis” pathways in the colonic lumen and mucosa samples were exclusively enriched in MUT compared to WT mice (**Figure 5C-E**, **Table S3-5**).

**Figure 5.**
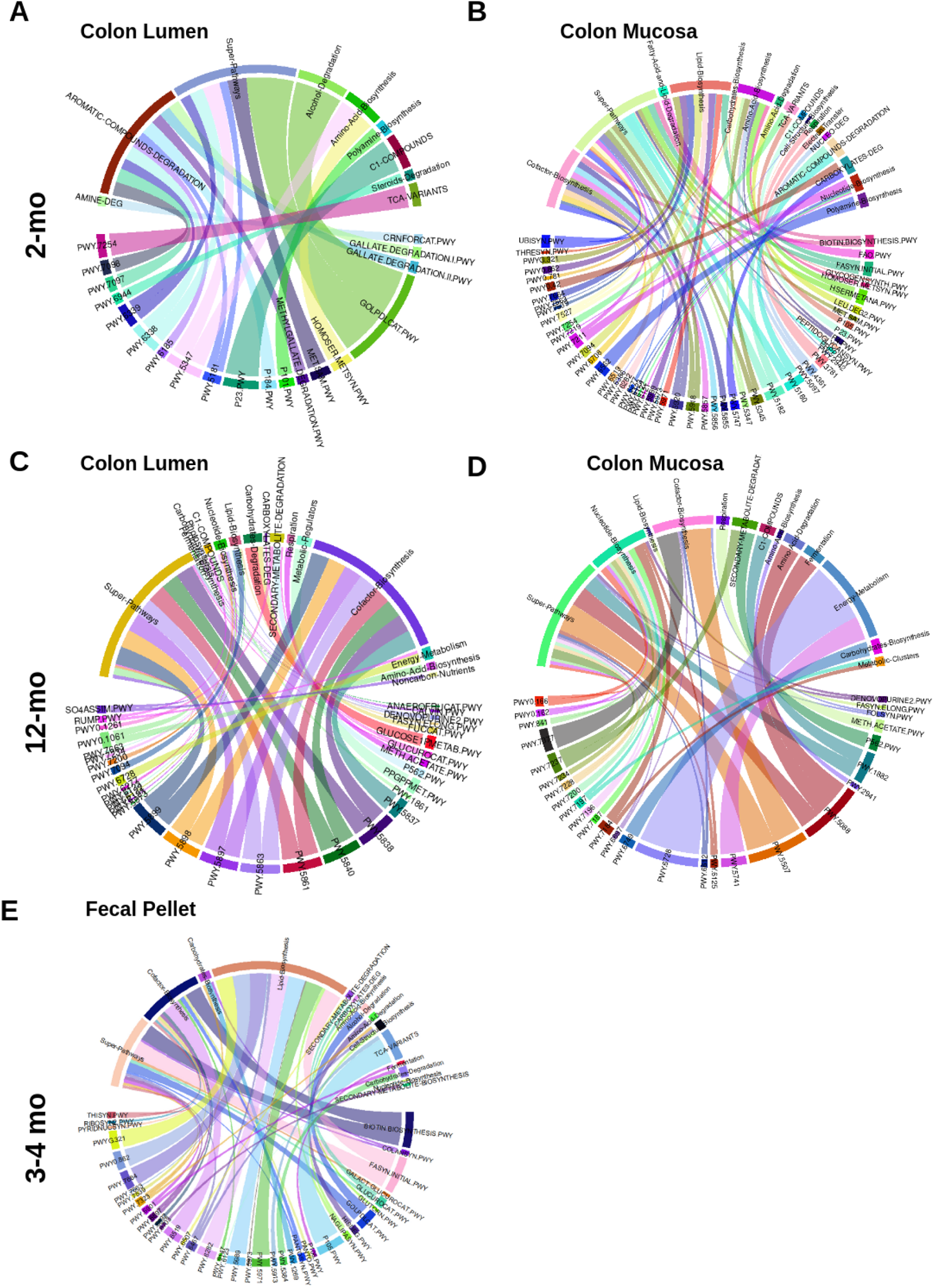
MUT mice exhibit differential abundance of predicted functional pathways. The PICRUSt2 algorithm was used to generate predicted MetaCyc pathway abundances from the 16S rRNA gene compositional data. Genotype-associated pathways were identified through linear mixed-effects regression. Circle plots group pathways which significantly distinguished (q<0.25) MUT from WT mice into higher-order categories, for the colonic luminal and colonic mucosal datasets of 2-month old (A and B) or 12-month old mice (C and D), or the fecal pellets of 3-4 month old mice (E).

### The SLC39A8 variant mediates spontaneous colonic inflammation but does not modulate susceptibility to chemically-induced colitis

To evaluate whether MUT mice exhibit intestinal inflammation in a manner consistent with the Crohn’s disease risk conferred by SLC39A8 A391T variant carriage in humans, we performed histological scoring of hematoxylin and eosin-stained colon tissue from 5-month old and 10-month old mice. There were no differences in inflammation between 5-month old MUT and WT mice (**Figure 6A-B**). Additionally, in contrast to what was previously reported, there were no differences in mucus barrier thickness as assessed through Alcian Blue/Periodic Acid Schiff staining between 6-month old MUT and WT mice (**Supplemental Figure 2**) ^9^. However, increased inflammation was observed in 10-month old MUT compared to WT mice (**Figure 6C-D**). This suggests that inflammation due to carriage of the Thr allele spontaneously develops, but may require an additional insult such as aging.

**Figure 6.**
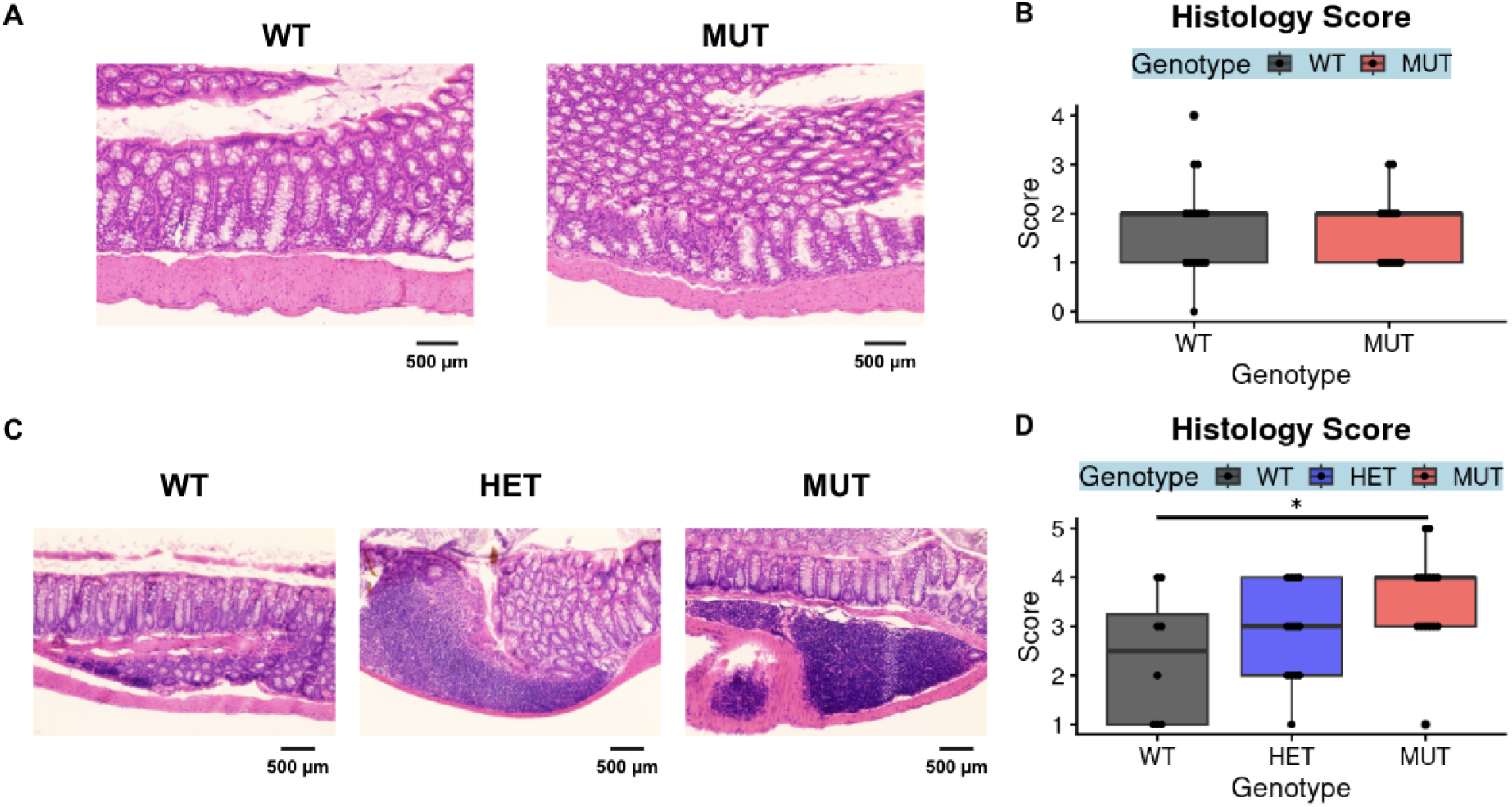
MUT mice exhibit age-dependent changes in spontaneous inflammations. Formalin-fixed, paraffin-embedded sections of WT and MUT colon tissue were stained with hematoxylin and eosin and scored for inflammation as assessed by lymphocyte infiltration and tissue architecture. Images and scores from colons from 5-month old mice (the ICP-MS cohort) are shown in (A, B), and images and scores from colons from 10-month old mice are shown in (C, D). Representative images were obtained through brightfield microscopy at 20x magnification (A, C) with a scale bar representing 500 microns. Significance was assessed through Wilcoxon rank-sum tests comparing WT to HET or WT to MUT, *p<0.05. The data in (A) and (B) represent 16 WT (7 F 9 M) and 13 MUT (4 F 9 M) mice, where the data in (C) and (D) represent n=12 WT (6 F 6 M), n=11 HET (5 F 6 M), and n=11 MUT (7 F 4 M) mice.

Two other studies have reported increased susceptibility of MUT compared to WT mice to dextran sodium sulfate (DSS)-induced colitis (one cohort aged 9-12 weeks, the other 11.5-13 weeks) ^9^ ^7^. We sought to reproduce these results by administering DSS to 9-12 week old WT, HET, and MUT mice to induce colitis. Mice were compared on three primary outcomes: body weight as a percent of baseline body weight prior to induction (where % baseline body weight decreases with increasing disease severity), colon length (where colon length is inversely related to inflammation), and colon histology score (where greater scores indicate higher disease severity) (**Figure 7**). Hematoxylin and eosin-stained colons from the DSS-treated cohort additionally underwent histological scoring. There were no differences by genotype in body weight, colon length, or histological scores for DSS-treated mice (**Figure 7A-D**). To confirm the absence of vulnerability to chemically induced colitis, we additionally utilized a second model - the 2,4,6-Trinitrobenzene sulfonic acid (TNBS) model. There were no differences by genotype in body weight or in colon length in the TNBS-treated mice (**Figure 7E-F**).

**Figure 7.**
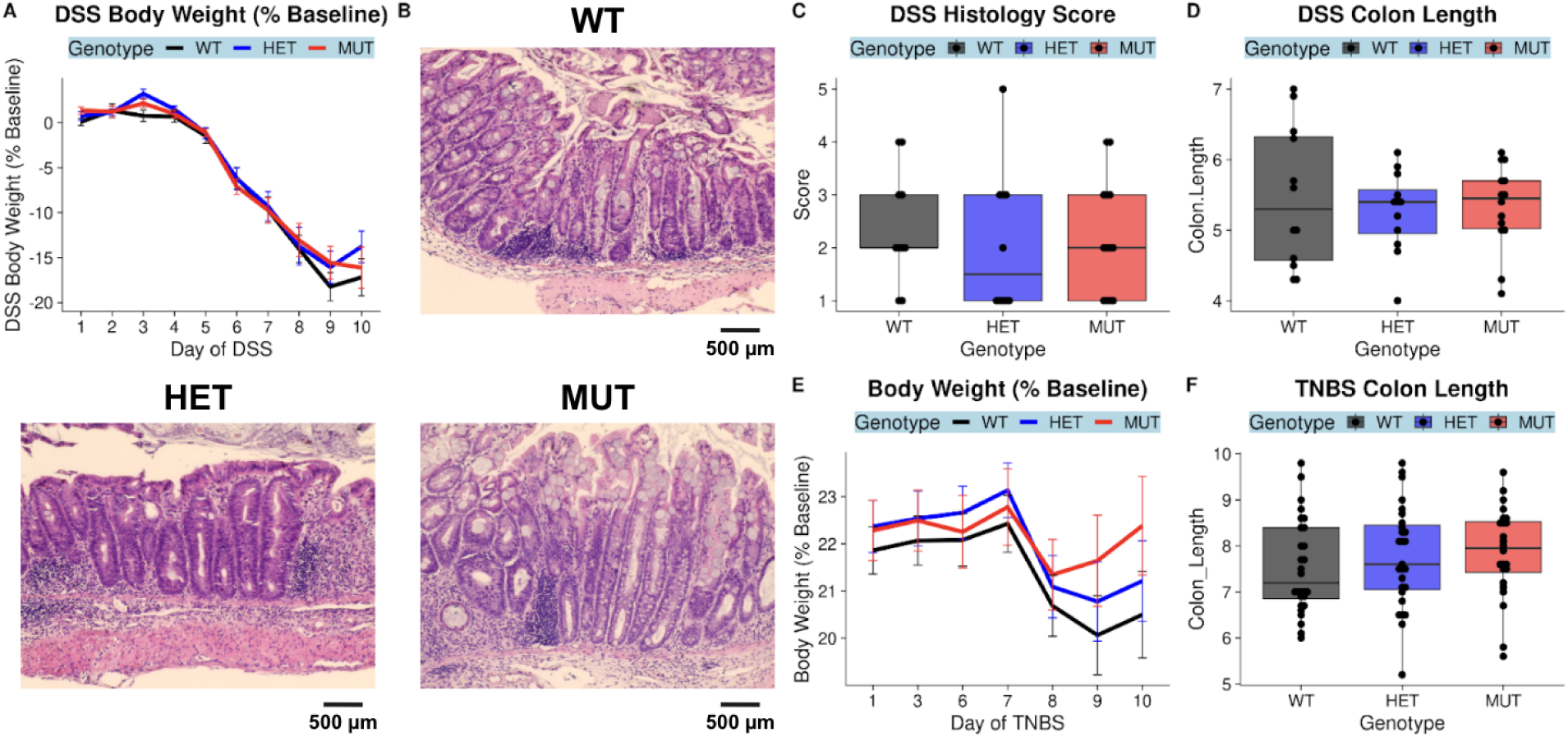
The A393T variant does not increase susceptibility to DSS- or TNBS-mediated chemical induction of colitis. DSS was administered via the drinking water to 2-3 month old mice, and body weight loss assessed as % of baseline was tracked for 10 days (A). Formalin-fixed, paraffin-embedded colon tissue sections were stained with hematoxylin and eosin and scored for inflammation. Representative colon images by genotype are shown in (B), and the distribution of histology scores by genotype are shown in (C). Differences in colon length (cm) by genotype are shown in (D). The DSS cohort represents n=13 WT mice (4 F 9 M), n=13 HET (4 F 9 M), and n=15 MUT (3 F 12 M) mice. In another cohort of 2-3 month old TNBS mice, mice were pre-sensitized on Day 0 and injected intrarectally with TNBS on Day 7. Body weight loss assessed as % of baseline was tracked for 3 days following the injection (E). Distributions of colon length (cm) by genotype for TNBS mice are shown in (F). Representative images were obtained through brightfield microscopy at 20x magnification (B) with a scale bar representing 500 microns. The TNBS cohort was comprised of n=20 WT (8 F 12 M), n=22 HET (8 F 14 M), and n=17 MUT (5 F 12 M) mice.

## DISCUSSION

The relationship between genetic risk, the human gut microbiome, and IBD is unclear but remains critical to enhancing predictions of disease risk and developing more personalized therapies. Emerging evidence from GWAS, including those with microbial abundances as quantitative traits ^21^, and microbiota studies in gene knockout preclinical models supports the notion that SNPs in susceptibility genes may confer disease risk through modulation of the microbiota ^22^ ^23^. Despite this, few studies have investigated the impact of specific risk loci on gut microbiome composition, aside from the Atg16L1 T300A variant^24^. We report here, for the first time, that the Crohn’s disease linked missense variant in SLC39A8 modulates colonic microbiome composition, consistent with what has previously been reported in human Crohn’s disease patients and in healthy control variant carriers ^6^. These results position SLC39A8 A391T as a possible Crohn’s disease-driving microbiome quantitative trait locus (mb-QTL).

Utilizing multiple cohorts of SLC39A8 A393T (matching human A391T) mice in this study, we demonstrated that the effects of A393T on microbiome community structure were apparent in 2-month old, 3-4 month old, and 1-year old mice. We additionally identified intestinal inflammation through histological analysis in 10-month old mice but not 5-month old mice. Though our study was not truly longitudinal, we can leverage this cross-sectional data from mice at different ages to conclude that A393T-linked microbiome dysbiosis occurs prior to the onset of intestinal inflammation. This supports the idea that risk variant carriage regulates the microbiome as a step in pathogenesis. These findings also mirror observations in a longitudinal study of Crohn’s disease patients, where higher instability in the gut microbiome in the remission phase was associated with higher risk of flare ^25^.

To better understand this A393T-linked microbiome signature on overall community structure, we performed differential taxon abundance testing and differential predicted pathway enrichment analysis in MUT compared to WT mice. While there were no concordant A393T-linked genera across different age cohorts, pathways mapping to “cofactor biosynthesis” were consistently differentially enriched in MUT compared to WT mice. Divalent cations transported by ZIP8, including iron, zinc, manganese, copper, and cobalt are utilized by members of the gut microbiome for processes including antioxidant response and metabolism ^13^ ^26^. These metal ions often serve as cofactors for enzymes involved in these processes, such as iron for heme ^27^. Increased exposure to several of these metals or metal-deficient diets have been shown to affect the composition of the gut microbiome ^28^ ^29^ ^30^ ^31^. At the colonic mucosal-luminal interface, we found that MUT mice exhibited reductions of iron, zinc, manganese, copper, cobalt, and cadmium in the lumen and increased abundance of specifically cobalt in the epithelial cells. Several of the predicted cofactor biosynthesis pathways differentially enriched in MUT mice employ metal-dependent enzymes. These included depletion of iron-dependent heme biosynthesis among mucosal-adherent bacteria of 2-month old mice, depletion of zinc-dependent folate biosynthesis in the feces of 3-4 month old mice, and enrichment of two cobalt pathways (cob(II)yrinate a,c-diamide biosynthesis I and adenosylcobalamin biosynthesis I) in the mucosal-adherent bacteria of 1-year old mice ^32^. Though the direction in which these predicted pathways are differentially enriched do not appear to concur with the trace element abundance results from ICP-MS, under conditions of metal depletion, some bacteria may have strategies to employ alternative metal-independent enzymes in critical processes ^32^. Thus, microbiome dysbiosis in MUT mice may be secondary to the metal dyshomeostasis in these mice. Future studies that include metatranscriptomics could provide further insight into bacterial responses to altered trace metal availability in MUT mice.

There were several notable differences between our findings and those of prior studies of SLC39A8 A393T mice. Consistent with a loss-of-function conferred by the variant, previous ICP-MS of A393T mice revealed that MUT compared to WT mice exhibited reduced Mn in blood and liver tissues ^9^ ^7^. However, while Sunuwar et al. found no differences in trace element abundance in the colons of MUT and WT mice aged 9-14 weeks of age, Nakata et al. reported 30-32 week old MUT mice exhibited reduced Mn abundance in MUT compared to WT colons. In bulk colon tissue, we actually found increased Mn abundance in MUT compared to WT colons. Our findings in 17-22 week old mice suggest that the consequences of the ZIP8 substitution at the apical surface of the colon is a gain-of function, where in MUT mice there is increased uptake of several metals while restricting the pool of available ions in the lumen. One possibility is that this discrepancy may be due to study-specific designs (e.g. age and sex) as well as experimental variability across studies with small sample size. Another possibility is that this is a physiologically-relevant outcome related to microbiome differences across facilities, revealing both strengths and weaknesses of performing these gene-microbiome studies. If the risk variant indeed confers disease susceptibility through its effects on the microbiome (which is additionally affected by environmental exposure), there is also the possibility that the microbes and microbe-derived signals affect the expression of other genes ^22^. With our facility-specific microbiome, this may manifest itself as upregulation of redundant transporters for Zn, Mn, and Fe, such as ZIP14, that compensate for reduced uptake by ZIP8 ^33^. Supporting this notion, Nakata et al. and Sunuwar et al. reported increased susceptibility to DSS-induced colitis in MUT compared to WT mice, while we did not observe any differences between the two. Nakata et al. also reported thinner mucus layers in MUT mice compared to WT mice that they proposed was due to a reduction in Mn-dependent glycosylases, but we did not observe a difference between MUT and WT mice. The importance of microbiome differences across facilities in the outcome of DSS colitis experiments has recently been demonstrated by Forster et al., who reported that susceptibility to DSS-induced colitis varied across facilities in a manner that depended on whether specific microbes were present within the microbiota of each facility ^34^.

In summary, we found that the SLC39A8 variant affects colonic mucosal-luminal trace metal homeostasis, changes the microbiome prior to disease onset, and promotes eventual spontaneous colitis representing a model for how it could promote disease in humans. Future SLC39A8 variant-associated microbiome transfer studies (from human and/or mouse variant carriers) will be critical to understanding whether the genetic risk-associated microbiome can indeed drive intestinal inflammation. Our study highlights the potential for other IBD-associated genetic risk variants to follow a similar paradigm in which the microbiome is altered as an intermediate step to the development of intestinal inflammation.

## DATA AVAILABILITY STATEMENT

Raw sequencing data is available on NCBI via the accession code PRJNA1004665 and will be made publicly available upon publication (reviewer link: https://dataview.ncbi.nlm.nih.gov/object/PRJNA1004665?reviewer=cmcbr110pvdd6pg1sotae449e9)

## AUTHOR CONTRIBUTIONS

Conceptualization: J.P.J. Data curation: J.C.Y. Formal analysis: J.C.Y. (Lead) M.Z. (Supporting) Funding acquisition: J.P.J. and D.C. Investigation: N.A.J., Z.Y., J.C.Y., (Lead) M.Z., D.C., J.S., A.G., C.C., F.L., V.L. (Supporting) Methodology: J.P.J., J.C.Y., V.L., N.A.J., Z.Y. Project administration: J.C.Y., N.A.J., Z.Y., V.L. Resources: J.P.J., V.L. Software: J.C.Y. Supervision: J.P.J., J.C.Y., N.A.J., Z.Y. Validation: M.Z. Visualization: J.C.Y., M.Z. Writing - original draft: J.C.Y. (Lead), M.Z., D.C. (Supporting). Writing - review & editing: J.P.J.

## ACKNOWLEDGEMENTS

The authors acknowledge the use of the ICP-MS facility within the UC Center for Environmental Implications of Nanotechnology in CNSI at UCLA, the Microbiome Core of the Goodman-Luskin Microbiome Center for performing microbiome sequencing, the UC Irvine Transgenic Mouse Facility for generation of the SLC38 A393T mice, and the Translational Pathology Core Laboratory for embedding, sectioning and performing the hematoxylin and eosin-staining of the intestines. The authors additionally acknowledge the use of BioRender.com in preparing the graphical abstract.

