## Supplemental Figures and Tables for "The ZIP8 A391T Crohn’s disease-linked risk variant induces colonic metal ion dyshomeostasis, microbiome compositional shifts, and inflammation"

**A**

```
      G  L  A  F  G  I  L  V  G  N  N  F  A  P  N  I  I  F  A  L  A  G  G  M  F  L  Y  I  S  L  A  D  M
WT    GGACTAGCTTTCGGCATTTTGGTGGGCAACAATTTTGCTCCCAATATTATATTTGCACTCGCTGGAGGCATGTTCCTCTACATTTCTCTGGCAGATATG

      G  L  A  F  G  I  L  V  G  N  N  F  T  P  N  I  I  F  A  L  A  G  G  M  F  L  Y  I  S  L  A  D  M
mut   GGACTAGCTTTCGGaATTcTcGTGGGCAACAATTTTacaCCCAAcATcATcTTTGCACTCGCTGGAGGCATGTTCCTCTACATTTCTCTGGCAGATATG

Proposed HDR sODN
GCTGTTCAACTTCCTCTCCGCGTGTTCCTGCTACGTGGGACTAGCTTTCGGaATTcTcGTGGGCAACAATTTTacaCCCAAcATcATcTTTGCACTCGCTGGAGGCA
TGTTCCTCTACATTTCTCTGGCAGATATGGTAAGAGA

sgRNA                 PAM
CTAGCTTTCGGCATTTTGGT  GGG
AGCGAGTGCAAATATAATAT  TGG
```

**B**

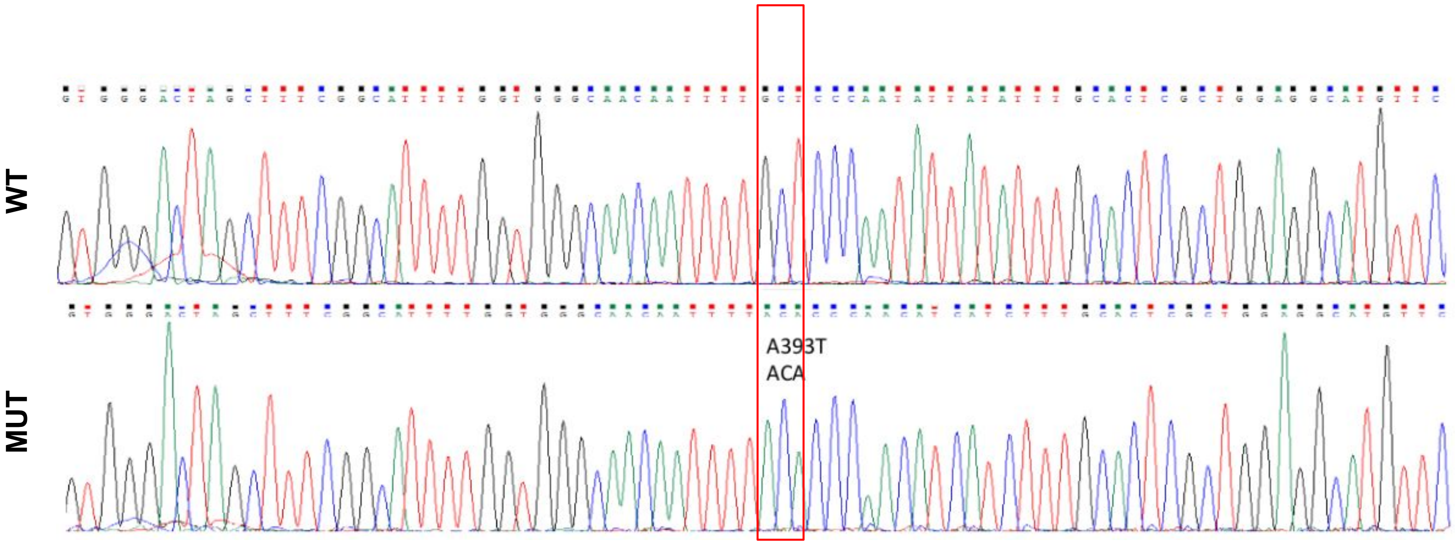


**Figure S1. Gene editing strategy to generate the SLC39A8 A393T knockin line.** (A) Nucleotide and corresponding amino acid sequence for the targeted sequence in SLC39A8 with **bolded and underlined** nucleotides corresponding to the 393 position. Regions highlighted yellow correspond to the regions targeted by the two guide RNA molecules (sgRNA). The single-stranded oligodeoxynucleotide sequence (sODN), used as the template for homology-directed recombination (HDR), contains the three point mutations used to generate the knockin along with additional silent mutations to prevent further Cas9-mediated mutation following HDR. The final gene-edited MUT mouse contains both the knockin and the silent mutations. (B) Sanger sequencing results demonstrate successful knockin in the gene-edited offspring.

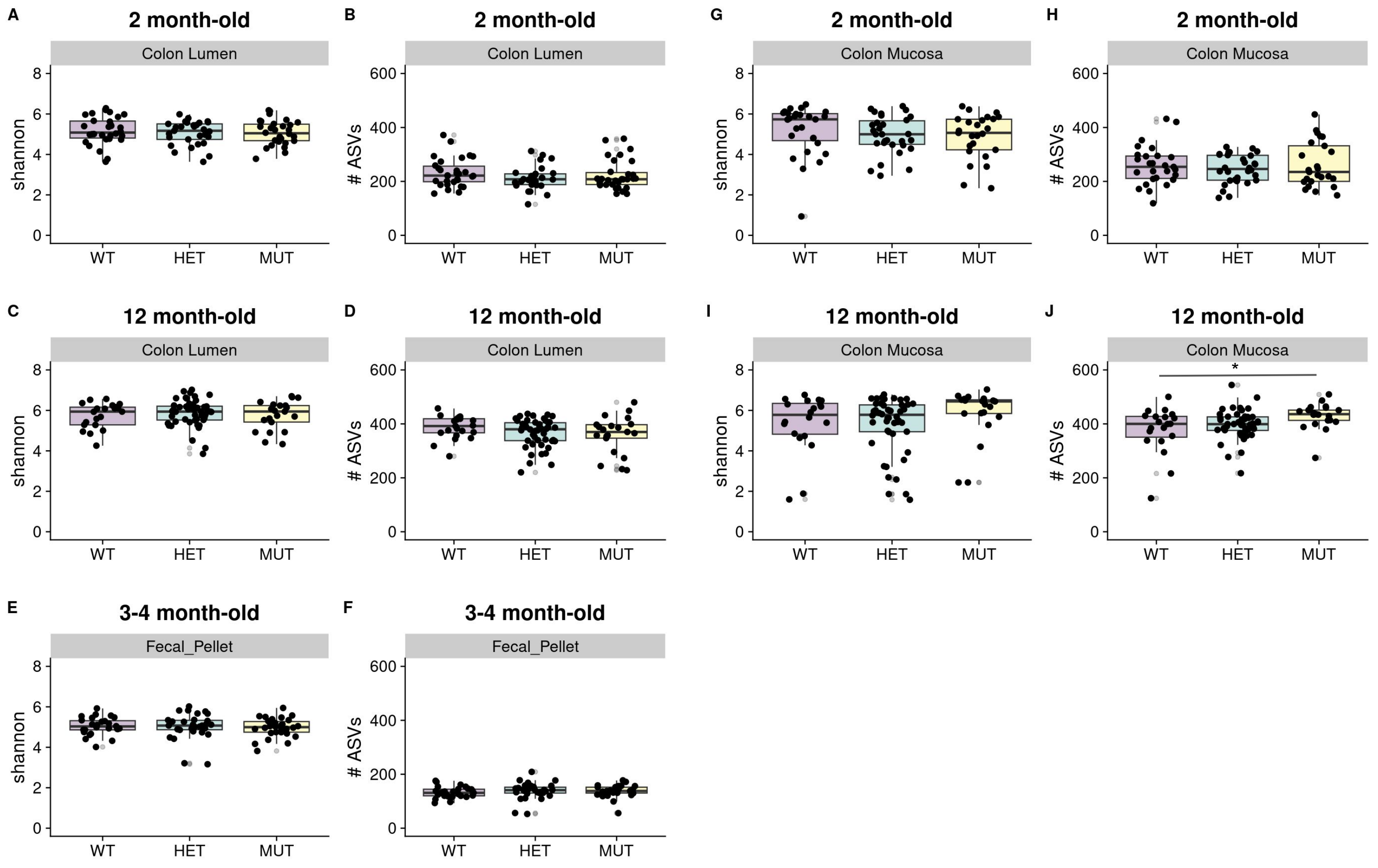


**Figure S2. Minimal differences in microbiome alpha-diversity are observed between genotypes.** Boxplots show the distribution of Shannon indices or the distribution of total number of amplicon sequence variants (# ASVs) across WT, HET, and MUT mice for colonic lumen datasets of 2 month-old (A, B), 12 month-old (C, D), or in the fecal pellets of 3-4 month-old mice (E, F). Distributions of Shannon index or of # ASVs are shown for the colonic mucosa datasets of 2 month-old (G, H) or 12 month-old mice (I, J). Significance of WT-HET, WT-MUT comparisons was assessed through linear mixed-effects models incorporating Sex and Genotype as fixed effects and MouseID as a random effect for the 2 month-old and 12 month-old datasets, or linear models with Sex and Genotype as covariates for the 3-4 months old dataset, *p<0.05. The data represents n=10 WT (4 F 6 M), n=10 HET (4 F 6 M), and n=10 MUT (4 F 6 M) for 2 month-old mice, and n=27 WT  (11 F 16 M), n=30 HET (10 F 20 M), and n=31 MUT (14 F 17 M) for 3-4 month-old mice.

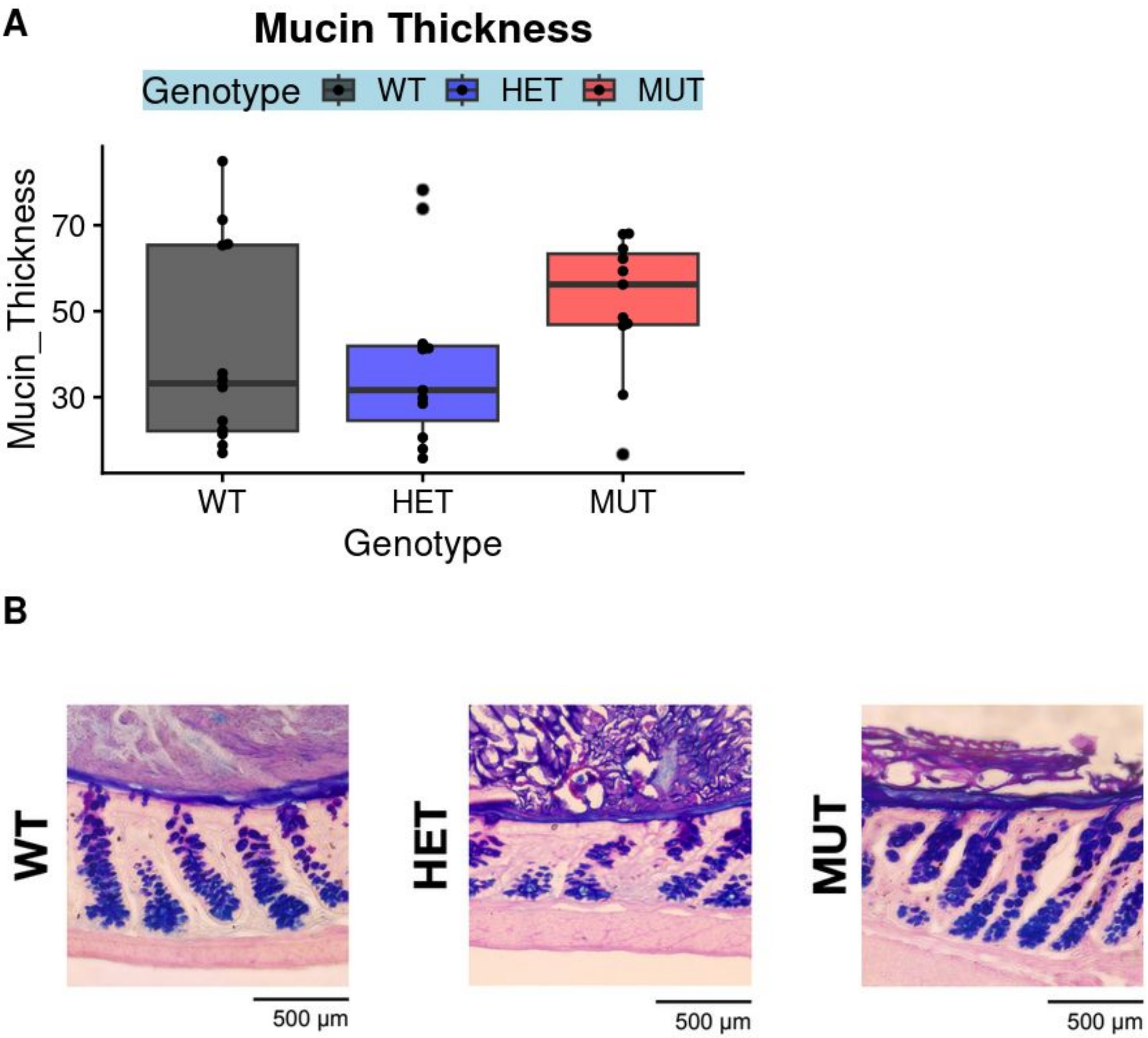


**Figure S3. MUT mice do not exhibit differential mucus layer thickness compared to WT mice.** Mucus layer staining of a separate cohort of 5-6 month old WT, HET, and MUT mice was achieved through PAS-Alcian Blue staining of paraffin-embedded, Carnoy's-fixed colon tissue. Mucus layer thickness measurements are shown in (A). Representative brightfield microscopy at 40X magnification of WT, HET, and MUT mucus layers are shown in (B). This data represents 5-6 month old n=12 WT (6 F 6 M), n=11 HET (5 F 6 M), and n=11 MUT (7 F 4 M) mice.

**TABLE S1: 2-MONTH LUMINAL**

| feature | coef | pval | qval | description | category |
|---|---|---|---|---|---|
| GOLPDLCAT.PWY | -0.2417127694 | 0.08617388403 | 0.2173051554 | superpathway of glycerol degradation to 1,3-propanediol | Super-Pathways |
| GOLPDLCAT.PWY | -0.2417127694 | 0.08617388403 | 0.2173051554 | superpathway of glycerol degradation to 1,3-propanediol | Alcohol-Degradation |
| P23.PWY | -0.1696159337 | 0.07092567432 | 0.186317704 | reductive TCA cycle I | C1-COMPOUNDS |
| HOMOSER.METSYN.PWY | -0.1002579161 | 0.08769143883 | 0.2200968291 | L-methionine biosynthesis I | Amino-Acid-Biosynthesis |
| MET.SAM.PWY | -0.09628676507 | 0.09189800736 | 0.2295186686 | superpathway of S-adenosyl-L-methionine biosynthesis | Super-Pathways |
| PWY.5347 | -0.0877745348 | 0.07780239477 | 0.199978779 | superpathway of L-methionine biosynthesis (transsulfuration) | Super-Pathways |
| PWY.5347 | -0.0877745348 | 0.07780239477 | 0.199978779 | superpathway of L-methionine biosynthesis (transsulfuration) | Amino-Acid-Biosynthesis |
| PWY.5181 | 0.04332487159 | 0.02978705199 | 0.08818706779 | toluene degradation III (aerobic) (via p-cresol) | AROMATIC-COMPOUNDS-DEGRADATION |
| PWY.5181 | 0.04332487159 | 0.02978705199 | 0.08818706779 | toluene degradation III (aerobic) (via p-cresol) | Super-Pathways |
| PWY.6185 | 0.04362684804 | 0.04520969298 | 0.1273406352 | 4-methylcatechol degradation (ortho cleavage) | AROMATIC-COMPOUNDS-DEGRADATION |
| CRNFORCAT.PWY | 0.04964005154 | 0.02341694094 | 0.07066898247 | creatinine degradation I | AMINE-DEG |
| PWY.6944 | 0.05206931007 | 0.002833523204 | 0.01342613331 | androstenedione degradation | Steroids-Degradation |
| PWY.6339 | 0.0542396036 | 0.004030040732 | 0.01738919703 | syringate degradation | Super-Pathways |
| PWY.6339 | 0.0542396036 | 0.004030040732 | 0.01738919703 | syringate degradation | AROMATIC-COMPOUNDS-DEGRADATION |
| METHYLGALLATE.DEGRADATION.PWY | 0.0564689511 | 0.003739501911 | 0.01645267031 | methylgallate degradation | AROMATIC-COMPOUNDS-DEGRADATION |
| GALLATE.DEGRADATION.I.PWY | 0.05656847812 | 0.003712348129 | 0.01640227016 | gallate degradation II | AROMATIC-COMPOUNDS-DEGRADATION |
| GALLATE.DEGRADATION.II.PWY | 0.05656847812 | 0.003712348129 | 0.01640227016 | gallate degradation I | AROMATIC-COMPOUNDS-DEGRADATION |
| P184.PWY | 0.07060550796 | 0.000373150021 | 0.002342873818 | protocatechuate degradation I (meta-cleavage pathway) | AROMATIC-COMPOUNDS-DEGRADATION |
| PWY.7098 | 0.07090143887 | 0.000348319907 | 0.002213836857 | vanillin and vanillate degradation II | AROMATIC-COMPOUNDS-DEGRADATION |
| PWY.6338 | 0.07155336862 | 0.000388341156 | 0.002408427722 | superpathway of vanillin and vanillate degradation | AROMATIC-COMPOUNDS-DEGRADATION |
| PWY.6338 | 0.07155336862 | 0.000388341156 | 0.002408427722 | superpathway of vanillin and vanillate degradation | Super-Pathways |
| PWY.7097 | 0.07155336862 | 0.000388341156 | 0.002408427722 | vanillin and vanillate degradation I | AROMATIC-COMPOUNDS-DEGRADATION |
| P101.PWY | 0.07262945299 | 0.000363305913 | 0.002288150288 | ectoine biosynthesis | Polyamine-Biosynthesis |
| PWY.7254 | 0.1137949781 | 0.01863060502 | 0.05812748765 | TCA cycle VII (acetate-producers) | TCA-VARIANTS |

TABLE S2: 2-MONTH MUCOSAL

| feature | coef | pval | qval | description | category |
|---|---|---|---|---|---|
| PWY.6562 | -0.3572907034 | 0.04338831714 | 0.1279711601 | norspermidine biosynthesis | Polyamine-Biosynthesis |
| PWY.5180 | -0.2454901222 | 0.07939533833 | 0.2020356375 | toluene degradation I (aerobic) (via o-cresol) | AROMATIC-COMPOUNDS-DEGRADATION |
| PWY.5180 | -0.2454901222 | 0.07939533833 | 0.2020356375 | toluene degradation I (aerobic) (via o-cresol) | Super-Pathways |
| PWY.5182 | -0.2454901222 | 0.07939533833 | 0.2020356375 | toluene degradation II (aerobic) (via 4-methylcatechol) | AROMATIC-COMPOUNDS-DEGRADATION |
| PWY.5182 | -0.2454901222 | 0.07939533833 | 0.2020356375 | toluene degradation II (aerobic) (via 4-methylcatechol) | Super-Pathways |
| FAO.PWY | -0.2315183883 | 0.05864457784 | 0.1626864114 | fatty acid β-oxidation I | Fatty-Acid-and-Lipid-Degradation |
| PWY.7094 | -0.2295337934 | 0.05122863473 | 0.145378558 | fatty acid salvage | Lipid-Biosynthesis |
| PWY.5747 | -0.2222063201 | 0.06712086811 | 0.1795590102 | 2-methylcitrate cycle II | CARBOXYLATES-DEG |
| PWY0.42 | -0.2164195962 | 0.07292118528 | 0.1909407595 | 2-methylcitrate cycle I | CARBOXYLATES-DEG |
| LEU.DEG2.PWY | -0.1926349302 | 0.06302355578 | 0.1709941436 | L-leucine degradation I | Amino-Acid-Degradation |
| HSERMETANA.PWY | -0.1863974991 | 0.007714058653 | 0.0285705876 | L-methionine biosynthesis III | Super-Pathways |
| HSERMETANA.PWY | -0.1863974991 | 0.007714058653 | 0.0285705876 | L-methionine biosynthesis III | Amino-Acid-Biosynthesis |
| PWY.7254 | -0.1825255872 | 0.06949279994 | 0.184261212 | TCA cycle VII (acetate-producers) | TCA-VARIANTS |
| FASYN.INITIAL.PWY | -0.182079508 | 0.07966793885 | 0.2022151001 | superpathway of fatty acid biosynthesis initiation (E. coli) | Super-Pathways |
| FASYN.INITIAL.PWY | -0.182079508 | 0.07966793885 | 0.2022151001 | superpathway of fatty acid biosynthesis initiation (E. coli) | Lipid-Biosynthesis |
| PWY.6282 | -0.1801554406 | 0.07847659866 | 0.2012949118 | palmitoleate biosynthesis I (from (5Z)-dodec-5-enoate) | Lipid-Biosynthesis |
| PWY0.862 | -0.1800522863 | 0.07850439508 | 0.2012949118 | (5Z)-dodec-5-enoate biosynthesis | Lipid-Biosynthesis |
| PWY.5989 | -0.1798270195 | 0.07856464518 | 0.2012949118 | stearate biosynthesis II (bacteria and plants) | Lipid-Biosynthesis |
| PWY.7664 | -0.1773358678 | 0.07719313923 | 0.1993918725 | oleate biosynthesis IV (anaerobic) | Lipid-Biosynthesis |
| PWY.5920 | -0.1761590811 | 0.04391663539 | 0.1289859221 | superpathway of heme biosynthesis from glycine | Super-Pathways |
| PWY.5920 | -0.1761590811 | 0.04391663539 | 0.1289859221 | superpathway of heme biosynthesis from glycine | Cofactor-Biosynthesis |
| PWYG.321 | -0.1756829287 | 0.07976160442 | 0.2022151001 | mycolate biosynthesis | Lipid-Biosynthesis |
| P105.PWY | -0.1680747992 | 0.0243160698 | 0.07831862974 | TCA cycle IV (2-oxoglutarate decarboxylase) | TCA-VARIANTS |
| UBISYN.PWY | -0.1674300301 | 0.08860575466 | 0.220228013 | superpathway of ubiquinol-8 biosynthesis (prokaryotic) | Super-Pathways |
| UBISYN.PWY | -0.1674300301 | 0.08860575466 | 0.220228013 | superpathway of ubiquinol-8 biosynthesis (prokaryotic) | Cofactor-Biosynthesis |
| PWY.5971 | -0.1674273311 | 0.06947478374 | 0.184261212 | palmitate biosynthesis II (bacteria and plants) | Lipid-Biosynthesis |
| PWY.5855 | -0.1673666099 | 0.08915078403 | 0.2202548782 | ubiquinol-7 biosynthesis (prokaryotic) | Cofactor-Biosynthesis |
| PWY.5856 | -0.1673666099 | 0.08915078403 | 0.2202548782 | ubiquinol-9 biosynthesis (prokaryotic) | Cofactor-Biosynthesis |
| PWY.5857 | -0.1673666099 | 0.08915078403 | 0.2202548782 | ubiquinol-10 biosynthesis (prokaryotic) | Cofactor-Biosynthesis |
| PWY.6708 | -0.1673666099 | 0.08915078403 | 0.2202548782 | ubiquinol-8 biosynthesis (prokaryotic) | Cofactor-Biosynthesis |
| P23.PWY | -0.1650890467 | 0.02053571982 | 0.06780662203 | reductive TCA cycle I | C1-COMPOUNDS |
| PWY.5918 | -0.1645397294 | 0.09081634786 | 0.2241061463 | superpathay of heme biosynthesis from glutamate | Cofactor-Biosynthesis |
| PWY.5918 | -0.1645397294 | 0.09081634786 | 0.2241061463 | superpathay of heme biosynthesis from glutamate | Super-Pathways |
| PWY.6519 | -0.1601092137 | 0.07543829173 | 0.1963078223 | 8-amino-7-oxononanoate biosynthesis I | Cofactor-Biosynthesis |
| PWY.5345 | -0.1532502485 | 0.05747562041 | 0.1598775944 | superpathway of L-methionine biosynthesis (by sulfhydrylation) | Amino-Acid-Biosynthesis |
| PWY.5345 | -0.1532502485 | 0.05747562041 | 0.1598775944 | superpathway of L-methionine biosynthesis (by sulfhydrylation) | Super-Pathways |
| PWY.3781 | -0.1520513943 | 0.04722216692 | 0.1356587559 | aerobic respiration I (cytochrome c) | Respiration |
| PWY.3781 | -0.1520513943 | 0.04722216692 | 0.1356587559 | aerobic respiration I (cytochrome c) | Electron-Transfer |
| BIOTIN.BIOSYNTHESIS.PWY | -0.1507405079 | 0.0746198649 | 0.1944190028 | biotin biosynthesis I | Cofactor-Biosynthesis |
| BIOTIN.BIOSYNTHESIS.PWY | -0.1507405079 | 0.0746198649 | 0.1944190028 | biotin biosynthesis I | Super-Pathways |
| PWY.4361 | -0.1251996515 | 0.04507548637 | 0.1314701686 | S-methyl-5-thio-α-D-ribose 1-phosphate degradation | Amino-Acid-Biosynthesis |

| | | | | | |
|---|---|---|---|---|---|
| PWY.4361 | -0.1251996515 | 0.04507548637 | 0.1314701686 | S-methyl-5-thio-α-D-ribose 1-phosphate degradation | NUCLEO-DEG |
| PWY.7527 | -0.1238785711 | 0.045070561 | 0.1314701686 | L-methionine salvage cycle III | Super-Pathways |
| PWY.7527 | -0.1238785711 | 0.045070561 | 0.1314701686 | L-methionine salvage cycle III | Amino-Acid-Biosynthesis |
| PWY.7211 | -0.117624966 | 0.0574798018 | 0.1598775944 | superpathway of pyrimidine deoxyribonucleotides de novo biosyn | Nucleotide-Biosynthesis |
| PWY.7211 | -0.117624966 | 0.0574798018 | 0.1598775944 | superpathway of pyrimidine deoxyribonucleotides de novo biosyn | Super-Pathways |
| PWY.7211 | -0.117624966 | 0.0574798018 | 0.1598775944 | superpathway of pyrimidine deoxyribonucleotides de novo biosyn | Nucleotide-Biosynthesis |
| MET.SAM.PWY | -0.1052499717 | 0.03768726316 | 0.1138751836 | superpathway of S-adenosyl-L-methionine biosynthesis | Super-Pathways |
| HOMOSER.METSYN.PWY | -0.1050116684 | 0.04572973411 | 0.132852915 | L-methionine biosynthesis I | Amino-Acid-Biosynthesis |
| PWY0.781 | -0.08419247893 | 0.01697810603 | 0.05713785684 | aspartate superpathway | Super-Pathways |
| PWY.5347 | -0.08327932348 | 0.04823468178 | 0.1376261301 | superpathway of L-methionine biosynthesis (transsulfuration) | Super-Pathways |
| PWY.5347 | -0.08327932348 | 0.04823468178 | 0.1376261301 | superpathway of L-methionine biosynthesis (transsulfuration) | Amino-Acid-Biosynthesis |
| PRPP.PWY | -0.07838320487 | 0.08325919669 | 0.2093943869 | superpathway of histidine, purine, and pyrimidine biosynthesis | Super-Pathways |
| PWY.7539 | -0.0756711268 | 0.09604189297 | 0.2339767694 | 6-hydroxymethyl-dihydropterin diphosphate biosynthesis III (Chla | Cofactor-Biosynthesis |
| P4.PWY | -0.06874926073 | 0.0468231943 | 0.1352526933 | superpathway of L-lysine, L-threonine and L-methionine biosynth | Super-Pathways |
| P4.PWY | -0.06874926073 | 0.0468231943 | 0.1352526933 | superpathway of L-lysine, L-threonine and L-methionine biosynth | Amino-Acid-Biosynthesis |
| PWY.7663 | 0.0241476503 | 0.0982359006 | 0.2373940061 | gondoate biosynthesis (anaerobic) | Lipid-Biosynthesis |
| PWY.2942 | 0.02594394824 | 0.07731874524 | 0.1994709644 | L-lysine biosynthesis III | Amino-Acid-Biosynthesis |
| PWY.5973 | 0.02711215681 | 0.08848627501 | 0.220228013 | cis-vaccenate biosynthesis | Lipid-Biosynthesis |
| PWY.5097 | 0.02836571282 | 0.07810945032 | 0.2010169677 | L-lysine biosynthesis VI | Amino-Acid-Biosynthesis |
| PWY.7219 | 0.02983416912 | 0.1008275238 | 0.2421055881 | adenosine ribonucleotides de novo biosynthesis | Nucleotide-Biosynthesis |
| PWY.6386 | 0.03225171785 | 0.1003516908 | 0.2416726498 | UDP-N-acetylmuramoyl-pentapeptide biosynthesis II (lysine-cont | Cell-Structure-Biosynthesis |
| PEPTIDOGLYCANSYN.PWY | 0.03297105883 | 0.1031997572 | 0.2459926108 | peptidoglycan biosynthesis I (meso-diaminopimelate containing) | Super-Pathways |
| PEPTIDOGLYCANSYN.PWY | 0.03297105883 | 0.1031997572 | 0.2459926108 | peptidoglycan biosynthesis I (meso-diaminopimelate containing) | Cell-Structure-Biosynthesis |
| THRESYN.PWY | 0.03540821719 | 0.06111162697 | 0.1666786203 | superpathway of L-threonine biosynthesis | Amino-Acid-Biosynthesis |
| THRESYN.PWY | 0.03540821719 | 0.06111162697 | 0.1666786203 | superpathway of L-threonine biosynthesis | Super-Pathways |
| GLYCOGENSYNTH.PWY | 0.03776749024 | 0.09336619797 | 0.2285186664 | glycogen biosynthesis I (from ADP-D-Glucose) | Carbohydrates-Biosynthesis |
| PWY.6123 | 0.03853258501 | 0.05305124528 | 0.1495404229 | inosine-5'-phosphate biosynthesis I | Nucleotide-Biosynthesis |
| PWY.6121 | 0.03882833633 | 0.04252960828 | 0.1259692206 | 5-aminoimidazole ribonucleotide biosynthesis I | Nucleotide-Biosynthesis |
| PWY.6122 | 0.04246155463 | 0.04699095556 | 0.1353648926 | 5-aminoimidazole ribonucleotide biosynthesis II | Nucleotide-Biosynthesis |
| PWY.6277 | 0.04246155463 | 0.04699095556 | 0.1353648926 | superpathway of 5-aminoimidazole ribonucleotide biosynthesis | Nucleotide-Biosynthesis |
| PWY.6277 | 0.04246155463 | 0.04699095556 | 0.1353648926 | superpathway of 5-aminoimidazole ribonucleotide biosynthesis | Super-Pathways |

TABLE S3: 12-MONTH LUMINAL

| feature | coef | pval | qval | description | category |
|---|---|---|---|---|---|
| PPGPPMET.PWY | -0.5508761421 | 0.01123100342 | 0.06348797464 | ppGpp biosynthesis | Metabolic-Regulators |
| PWY0.1261 | -0.1459570701 | 0.0357194808 | 0.1609823176 | anhydromuropeptides recycling | SECONDARY-METABOLITE-DEGRADATION |
| PWY.7234 | -0.1088683543 | 0.04312955762 | 0.1796154995 | inosine-5'-phosphate biosynthesis III | Nucleotide-Biosynthesis |
| PWY.7200 | -0.0869414218 | 0.06706238976 | 0.2490239981 | superpathway of pyrimidine deoxyribonucleoside salvage | Nucleotide-Biosynthesis |
| PWY.7200 | -0.0869414218 | 0.06706238976 | 0.2490239981 | superpathway of pyrimidine deoxyribonucleoside salvage | Super-Pathways |
| FASYN.ELONG.PWY | -0.08148658986 | 0.02652394511 | 0.1272066755 | fatty acid elongation -- saturated | Lipid-Biosynthesis |
| DENOVOPURINE2.PWY | -0.08093870563 | 0.0618559005 | 0.2366347822 | superpathway of purine nucleotides de novo biosynthesis II | Super-Pathways |
| DENOVOPURINE2.PWY | -0.08093870563 | 0.0618559005 | 0.2366347822 | superpathway of purine nucleotides de novo biosynthesis II | Nucleotide-Biosynthesis |
| PWY.6122 | -0.0484202216 | 0.014599104 | 0.07925916199 | 5-aminoimidazole ribonucleotide biosynthesis II | Nucleotide-Biosynthesis |
| PWY.6277 | -0.0484202216 | 0.014599104 | 0.07925916199 | superpathway of 5-aminoimidazole ribonucleotide biosynthesis | Super-Pathways |
| PWY.6277 | -0.0484202216 | 0.014599104 | 0.07925916199 | superpathway of 5-aminoimidazole ribonucleotide biosynthesis | Nucleotide-Biosynthesis |
| PWY.6121 | -0.04824193883 | 0.009817611291 | 0.05607628671 | 5-aminoimidazole ribonucleotide biosynthesis I | Nucleotide-Biosynthesis |
| PWY.6123 | -0.04283016331 | 0.007582197311 | 0.04535532573 | inosine-5'-phosphate biosynthesis I | Nucleotide-Biosynthesis |
| PWY.5973 | -0.0368747843 | 0.02145622999 | 0.1108581466 | cis-vaccenate biosynthesis | Lipid-Biosynthesis |
| PWY.7663 | -0.03145243878 | 0.04225453541 | 0.1768669485 | gondoate biosynthesis (anaerobic) | Lipid-Biosynthesis |
| PWY0.1061 | 0.2902366852 | 0.00392728895 | 0.02587997226 | superpathway of L-alanine biosynthesis | Amino-Acid-Biosynthesis |
| CALVIN.PWY | 0.03313705214 | 0.06201226606 | 0.2366825468 | Calvin-Benson-Bassham cycle | C1-COMPOUNDS |
| PWY.1861 | 0.1296586209 | 0.0196816249 | 0.1031091496 | formaldehyde assimilation II (RuMP Cycle) | C1-COMPOUNDS |
| RUMP.PWY | 0.1168975016 | 0.01613646413 | 0.0870310934 | formaldehyde oxidation I | C1-COMPOUNDS |
| CALVIN.PWY | 0.03313705214 | 0.06201226606 | 0.2366825468 | Calvin-Benson-Bassham cycle | Carbohydrates-Biosynthesis |
| FUCCAT.PWY | 0.195964064 | 0.02148649504 | 0.1108581466 | fucose degradation | Carbohydrates-Degradation |
| GLUCOSE1PMETAB.PWY | 0.4829588018 | 0.02784290565 | 0.1323745081 | glucose and glucose-1-phosphate degradation | Carbohydrates-Degradation |
| GLUCUROCAT.PWY | 0.07161366127 | 0.05788509488 | 0.2235289734 | superpathway of β-D-glucuronide and D-glucuronate degradation | CARBOXYLATES-DEG |
| PWY.5837 | 0.7259970898 | 0.008380980788 | 0.04866923167 | 1,4-dihydroxy-2-naphthoate biosynthesis I | Cofactor-Biosynthesis |
| PWY.5861 | 0.7106362482 | 0.008101471582 | 0.04742676425 | superpathway of demethylmenaquinol-8 biosynthesis | Cofactor-Biosynthesis |
| PWY.5897 | 0.702184886 | 0.008028673755 | 0.04716845831 | superpathway of menaquinol-11 biosynthesis | Cofactor-Biosynthesis |
| PWY.5898 | 0.702184886 | 0.008028673755 | 0.04716845831 | superpathway of menaquinol-12 biosynthesis | Cofactor-Biosynthesis |
| PWY.5899 | 0.702184886 | 0.008028673755 | 0.04716845831 | superpathway of menaquinol-13 biosynthesis | Cofactor-Biosynthesis |
| PWY.5840 | 0.6977595672 | 0.007948710223 | 0.04716845831 | superpathway of menaquinol-7 biosynthesis | Cofactor-Biosynthesis |
| PWY.5838 | 0.6956045908 | 0.007906246227 | 0.04712237335 | superpathway of menaquinol-8 biosynthesis I | Cofactor-Biosynthesis |
| PWY.5863 | 0.7248700561 | 0.008346371978 | 0.04866923167 | superpathway of phylloquinol biosynthesis | Cofactor-Biosynthesis |
| RUMP.PWY | 0.1168975016 | 0.01613646413 | 0.0870310934 | formaldehyde oxidation I | Energy-Metabolism |
| PWY.6728 | 0.3453100062 | 0.03913292048 | 0.1715769693 | methylaspartate cycle | Energy-Metabolism |
| ANAEROFRUCAT.PWY | 0.02207362847 | 0.03969342319 | 0.1720825046 | homolactic fermentation | Fermentation |
| PWY.7094 | 0.2652796396 | 0.01459166715 | 0.07925916199 | fatty acid salvage | Lipid-Biosynthesis |
| SO4ASSIM.PWY | 0.1423507482 | 0.06150767 | 0.2358510889 | sulfate reduction I (assimilatory) | Noncarbon-Nutrients |
| CALVIN.PWY | 0.03313705214 | 0.06201226606 | 0.2366825468 | Calvin-Benson-Bassham cycle | Photosynthesis |
| METH.ACETATE.PWY | 0.2589119533 | 0.02385397397 | 0.118908446 | methanogenesis from acetate | Respiration |
| P562.PWY | 0.2988172388 | 0.03165112424 | 0.1470793767 | myo-inositol degradation I | SECONDARY-METABOLITE-DEGRADATION |
| GLUCUROCAT.PWY | 0.07161366127 | 0.05788509488 | 0.2235289734 | superpathway of β-D-glucuronide and D-glucuronate degradation | SECONDARY-METABOLITE-DEGRADATION |

| | | | | | |
|---|---|---|---|---|---|
| ANAEROFRUCAT.PWY | 0.02207362847 | 0.03969342319 | 0.1720825046 | homolactic fermentation | Super-Pathways |
| SO4ASSIM.PWY | 0.1423507482 | 0.06150767 | 0.2358510889 | sulfate reduction I (assimilatory) | Super-Pathways |
| GLUCUROCAT.PWY | 0.07161366127 | 0.05788509488 | 0.2235289734 | superpathway of β-D-glucuronide and D-glucuronate degradation | Super-Pathways |
| PWY.5861 | 0.7106362482 | 0.008101471582 | 0.04742676425 | superpathway of demethylmenaquinol-8 biosynthesis | Super-Pathways |
| PWY0.1061 | 0.2902366852 | 0.00392728895 | 0.02587997226 | superpathway of L-alanine biosynthesis | Super-Pathways |
| PWY.5897 | 0.702184886 | 0.008028673755 | 0.04716845831 | superpathway of menaquinol-11 biosynthesis | Super-Pathways |
| PWY.5898 | 0.702184886 | 0.008028673755 | 0.04716845831 | superpathway of menaquinol-12 biosynthesis | Super-Pathways |
| PWY.5899 | 0.702184886 | 0.008028673755 | 0.04716845831 | superpathway of menaquinol-13 biosynthesis | Super-Pathways |
| PWY.5840 | 0.6977595672 | 0.007948710223 | 0.04716845831 | superpathway of menaquinol-7 biosynthesis | Super-Pathways |
| PWY.5838 | 0.6956045908 | 0.007906246227 | 0.04712237335 | superpathway of menaquinol-8 biosynthesis I | Super-Pathways |
| PWY.5863 | 0.7248700561 | 0.008346371978 | 0.04866923167 | superpathway of phylloquinol biosynthesis | Super-Pathways |

| TABLE S4: 12-MONTH MUCOSAL | | | | | |
|---|---|---|---|---|---|
| feature | coef | pval | qval | description | category |
| PWY.7234 | -0.1197205064 | 0.02613123826 | 0.07537857191 | inosine-5'-phosphate biosynthesis III | Nucleotide-Biosynthesis |
| PWY.7184 | -0.1044344694 | 0.05774941942 | 0.1573839555 | pyrimidine deoxyribonucleotides de novo biosynthesis I | Nucleotide-Biosynthesis |
| PWY.7184 | -0.1044344694 | 0.05774941942 | 0.1573839555 | pyrimidine deoxyribonucleotides de novo biosynthesis I | Nucleotide-Biosynthesis |
| PWY.7184 | -0.1044344694 | 0.05774941942 | 0.1573839555 | pyrimidine deoxyribonucleotides de novo biosynthesis I | Metabolic-Clusters |
| PWY.2941 | -0.1032063947 | 0.04875851293 | 0.1352399582 | L-lysine biosynthesis II | Amino-Acid-Biosynthesis |
| PWY.7197 | -0.1014718125 | 0.06272105842 | 0.1696998334 | pyrimidine deoxyribonucleotide phosphorylation | Nucleotide-Biosynthesis |
| PWY.7197 | -0.1014718125 | 0.06272105842 | 0.1696998334 | pyrimidine deoxyribonucleotide phosphorylation | Metabolic-Clusters |
| PWY.7196 | -0.08804402921 | 0.06122986707 | 0.1661447189 | superpathway of pyrimidine ribonucleosides salvage | Nucleotide-Biosynthesis |
| PWY.7196 | -0.08804402921 | 0.06122986707 | 0.1661447189 | superpathway of pyrimidine ribonucleosides salvage | Super-Pathways |
| PWY.7228 | -0.08764367227 | 0.07218540504 | 0.1936303783 | superpathway of guanosine nucleotides de novo biosynthesis I | Super-Pathways |
| PWY.7228 | -0.08764367227 | 0.07218540504 | 0.1936303783 | superpathway of guanosine nucleotides de novo biosynthesis I | Nucleotide-Biosynthesis |
| PWY.6125 | -0.08139965848 | 0.07297719314 | 0.1954746245 | superpathway of guanosine nucleotides de novo biosynthesis II | Super-Pathways |
| PWY.6125 | -0.08139965848 | 0.07297719314 | 0.1954746245 | superpathway of guanosine nucleotides de novo biosynthesis II | Nucleotide-Biosynthesis |
| PWY.7200 | -0.07755808087 | 0.07406579578 | 0.1978253093 | superpathway of pyrimidine deoxyribonucleoside salvage | Nucleotide-Biosynthesis |
| PWY.7200 | -0.07755808087 | 0.07406579578 | 0.1978253093 | superpathway of pyrimidine deoxyribonucleoside salvage | Super-Pathways |
| PWY0.162 | -0.07624519076 | 0.05802992895 | 0.1579188923 | superpathway of pyrimidine ribonucleotides de novo biosynthesis | Nucleotide-Biosynthesis |
| PWY0.162 | -0.07624519076 | 0.05802992895 | 0.1579188923 | superpathway of pyrimidine ribonucleotides de novo biosynthesis | Super-Pathways |
| PWY0.166 | -0.07503903496 | 0.09701472218 | 0.2488407717 | superpathway of pyrimidine deoxyribonucleotides de novo biosynthesis (E. coli) | Nucleotide-Biosynthesis |
| PWY0.166 | -0.07503903496 | 0.09701472218 | 0.2488407717 | superpathway of pyrimidine deoxyribonucleotides de novo biosynthesis (E. coli) | Super-Pathways |
| PWY0.166 | -0.07503903496 | 0.09701472218 | 0.2488407717 | superpathway of pyrimidine deoxyribonucleotides de novo biosynthesis (E. coli) | Nucleotide-Biosynthesis |
| PWY.7187 | -0.07496492223 | 0.08732852726 | 0.2277343374 | pyrimidine deoxyribonucleotides de novo biosynthesis II | Nucleotide-Biosynthesis |
| PWY.7187 | -0.07496492223 | 0.08732852726 | 0.2277343374 | pyrimidine deoxyribonucleotides de novo biosynthesis II | Nucleotide-Biosynthesis |
| DENOVOPURINE2.PWY | -0.07070018531 | 0.05643161779 | 0.1542409378 | superpathway of purine nucleotides de novo biosynthesis II | Super-Pathways |
| DENOVOPURINE2.PWY | -0.07070018531 | 0.05643161779 | 0.1542409378 | superpathway of purine nucleotides de novo biosynthesis II | Nucleotide-Biosynthesis |
| PWY.841 | -0.07042416458 | 0.05917639493 | 0.160805421 | superpathway of purine nucleotides de novo biosynthesis I | Nucleotide-Biosynthesis |
| PWY.841 | -0.07042416458 | 0.05917639493 | 0.160805421 | superpathway of purine nucleotides de novo biosynthesis I | Super-Pathways |
| FASYN.ELONG.PWY | -0.07018233925 | 0.04863063751 | 0.1350851042 | fatty acid elongation -- saturated | Lipid-Biosynthesis |
| PWY.5088 | 0.4500994296 | 0.02901318937 | 0.0831800154 | L-glutamate degradation VIII (to propanoate) | Amino-Acid-Degradation |
| PWY.1882 | 0.3392060786 | 0.08320947032 | 0.2175979872 | superpathway of C1 compounds oxidation to CO2 | C1-COMPOUNDS |
| PWY.6749 | 0.3259263014 | 0.04440767684 | 0.1235376767 | CMP-legionaminate biosynthesis I | Carbohydrates-Biosynthesis |
| PWY.6897 | 0.04865136887 | 0.07893880935 | 0.2084880187 | thiamin salvage II | Cofactor-Biosynthesis |
| FOLSYN.PWY | 0.05245942723 | 0.02413390877 | 0.07004810982 | superpathway of tetrahydrofolate biosynthesis and salvage | Cofactor-Biosynthesis |
| PWY.6612 | 0.0691851483 | 0.04242079623 | 0.1188923661 | superpathway of tetrahydrofolate biosynthesis | Cofactor-Biosynthesis |
| PWY.7377 | 0.4925770223 | 0.09455151804 | 0.2435221103 | cob(II)yrinate a,c-diamide biosynthesis I (early cobalt insertion) | Cofactor-Biosynthesis |
| PWY.5507 | 0.6496178914 | 0.02358443438 | 0.06855940227 | adenosylcobalamin biosynthesis I (early cobalt insertion) | Cofactor-Biosynthesis |
| PWY.5741 | 0.5453314262 | 0.008032035721 | 0.02448791378 | ethylmalonyl-CoA pathway | Energy-Metabolism |
| PWY.6728 | 1.374024239 | 0.001335409665 | 0.00436218314 | methylaspartate cycle | Energy-Metabolism |
| PWY.5088 | 0.4500994296 | 0.02901318937 | 0.0831800154 | L-glutamate degradation VIII (to propanoate) | Fermentation |
| METH.ACETATE.PWY | 0.2784705787 | 0.08843731799 | 0.2293498911 | methanogenesis from acetate | Respiration |
| PWY.7237 | 0.2137000538 | 0.04331254576 | 0.1212104825 | myo-, chiro- and scillo-inositol degradation | SECONDARY-METABOLITE-DEGRADATION |

| | | | | | |
|---|---|---|---|---|---|
| P562.PWY | 0.3332018527 | 0.01677014965 | 0.04983206115 | myo-inositol degradation I | SECONDARY-METABOLITE-DEGRADATION |
| PWY.6897 | 0.04865136887 | 0.07893880935 | 0.2084880187 | thiamin salvage II | Super-Pathways |
| FOLSYN.PWY | 0.05245942723 | 0.02413390877 | 0.07004810982 | superpathway of tetrahydrofolate biosynthesis and salvage | Super-Pathways |
| PWY.6612 | 0.0691851483 | 0.04242079623 | 0.1188923661 | superpathway of tetrahydrofolate biosynthesis | Super-Pathways |
| PWY.7237 | 0.2137000538 | 0.04331254576 | 0.1212104825 | myo-, chiro- and scillo-inositol degradation | Super-Pathways |
| PWY.1882 | 0.3392060786 | 0.08320947032 | 0.2175979872 | superpathway of C1 compounds oxidation to CO2 | Super-Pathways |
| PWY.5088 | 0.4500994296 | 0.02901318937 | 0.0831800154 | L-glutamate degradation VIII (to propanoate) | Super-Pathways |
| PWY.5507 | 0.6496178914 | 0.02358443438 | 0.06855940227 | adenosylcobalamin biosynthesis I (early cobalt insertion) | Super-Pathways |

| TABLE S5: 3-4 MONTH FECAL PELLETS | | | | | |
|---|---|---|---|---|---|
| feature | coef | pval | qval | description | category |
| GOLPDLCAT.PWY | -0.1769427398 | 0.0149836236 | 0.1756881841 | superpathway of glycerol degradation to 1,3-propanediol | Alcohol-Degradation |
| GLUTORN.PWY | -0.04645535177 | 0.04191150945 | 0.2496335161 | L-ornithine biosynthesis | Amino-Acid-Biosynthesis |
| COLANSYN.PWY | -0.04415063059 | 0.01751043647 | 0.1756881841 | colanic acid building blocks biosynthesis | Carbohydrates-Biosynthesis |
| PWY.1269 | -0.06044474317 | 0.03545778741 | 0.2290113368 | CMP-3-deoxy-D-manno-octulosonate biosynthesis I | Carbohydrates-Biosynthesis |
| PWY.7323 | -0.04961117381 | 0.01961060837 | 0.1756881841 | superpathway of GDP-mannose-derived O-antigen building blocks biosynthesis | Carbohydrates-Biosynthesis |
| PWY.1269 | -0.06044474317 | 0.03545778741 | 0.2290113368 | CMP-3-deoxy-D-manno-octulosonate biosynthesis I | Carbohydrates-Biosynthesis |
| PWY.6901 | -0.06323344154 | 0.02041669144 | 0.1756881841 | superpathway of glucose and xylose degradation | Carbohydrates-Degradation |
| PWY.5384 | -0.1318414954 | 0.009895895383 | 0.141667555 | sucrose degradation IV (sucrose phosphorylase) | Carbohydrates-Degradation |
| GLUCUROCAT.PWY | -0.05970682386 | 0.005041720407 | 0.09700305131 | superpathway of β-D-glucuronide and D-glucuronate degradation | CARBOXYLATES-DEG |
| NAGLIPASYN.PWY | -0.06151006075 | 0.03159354502 | 0.2130606011 | lipid IVA biosynthesis | Cell-Structure-Biosynthesis |
| PWY.6467 | -0.06110900162 | 0.03241046356 | 0.2150157583 | Kdo transfer to lipid IVA III (Chlamydia) | Cell-Structure-Biosynthesis |
| PWY.7323 | -0.04961117381 | 0.01961060837 | 0.1756881841 | superpathway of GDP-mannose-derived O-antigen building blocks biosynthesis | Cell-Structure-Biosynthesis |
| PANTOSYN.PWY | -0.04086930306 | 0.004019143793 | 0.09595114474 | pantothenate and coenzyme A biosynthesis I | Cofactor-Biosynthesis |
| PANTOSYN.PWY | -0.04086930306 | 0.004019143793 | 0.09595114474 | pantothenate and coenzyme A biosynthesis I | Cofactor-Biosynthesis |
| PWY.6897 | -0.06270223199 | 0.001512684015 | 0.0726088327 | thiamin salvage II | Cofactor-Biosynthesis |
| PWY.7539 | -0.05668053461 | 0.0051116804 | 0.09700305131 | 6-hydroxymethyl-dihydropterin diphosphate biosynthesis III (Chlamydia) | Cofactor-Biosynthesis |
| PWY.6147 | -0.05117317926 | 0.02120566006 | 0.1783898825 | 6-hydroxymethyl-dihydropterin diphosphate biosynthesis I | Cofactor-Biosynthesis |
| PYRIDNUCSYN.PWY | -0.03947295541 | 0.01470717029 | 0.1756881841 | NAD biosynthesis I (from aspartate) | Cofactor-Biosynthesis |
| PWY.6892 | -0.05862628088 | 0.00211127126 | 0.08203796894 | thiazole biosynthesis I (E. coli) | Cofactor-Biosynthesis |
| THISYN.PWY | -0.05608300397 | 0.009683246067 | 0.1410987284 | superpathway of thiamin diphosphate biosynthesis I | Cofactor-Biosynthesis |
| PANTO.PWY | -0.05040195979 | 0.01264524329 | 0.1657219763 | phosphopantothenate biosynthesis I | Cofactor-Biosynthesis |
| RIBOSYN2.PWY | -0.04491734943 | 0.000982103555 | 0.05161070219 | flavin biosynthesis I (bacteria and plants) | Cofactor-Biosynthesis |
| P108.PWY | -0.05611345943 | 0.01687656697 | 0.1756881841 | pyruvate fermentation to propanoate I | Fermentation |
| NAGLIPASYN.PWY | -0.06151006075 | 0.03159354502 | 0.2130606011 | lipid IVA biosynthesis | Lipid-Biosynthesis |
| PWY.6467 | -0.06110900162 | 0.03241046356 | 0.2150157583 | Kdo transfer to lipid IVA III (Chlamydia) | Lipid-Biosynthesis |
| PWY.6703 | -0.05637295488 | 0.02631912634 | 0.1983177847 | preQ0 biosynthesis | SECONDARY-METABOLITE-BIOSYNTHESIS |
| PWY.6507 | -0.04562301245 | 0.02790938226 | 0.1983177847 | 4-deoxy-L-threo-hex-4-enopyranuronate degradation | SECONDARY-METABOLITE-DEGRADATION |
| GLUCUROCAT.PWY | -0.05970682386 | 0.005041720407 | 0.09700305131 | superpathway of β-D-glucuronide and D-glucuronate degradation | SECONDARY-METABOLITE-DEGRADATION |
| GOLPDLCAT.PWY | -0.1769427398 | 0.0149836236 | 0.1756881841 | superpathway of glycerol degradation to 1,3-propanediol | Super-Pathways |
| PWY.6901 | -0.06323344154 | 0.02041669144 | 0.1756881841 | superpathway of glucose and xylose degradation | Super-Pathways |
| PWY.6897 | -0.06270223199 | 0.001512684015 | 0.0726088327 | thiamin salvage II | Super-Pathways |
| PWY.6467 | -0.06110900162 | 0.03241046356 | 0.2150157583 | Kdo transfer to lipid IVA III (Chlamydia) | Super-Pathways |
| GLUCUROCAT.PWY | -0.05970682386 | 0.005041720407 | 0.09700305131 | superpathway of β-D-glucuronide and D-glucuronate degradation | Super-Pathways |
| THISYN.PWY | -0.05608300397 | 0.009683246067 | 0.1410987284 | superpathway of thiamin diphosphate biosynthesis I | Super-Pathways |
| GALACT.GLUCUROCAT.PWY | -0.05352623777 | 0.01389390655 | 0.1756881841 | superpathway of hexuronide and hexuronate degradation | Super-Pathways |
| PWY.7323 | -0.04961117381 | 0.01961060837 | 0.1756881841 | superpathway of GDP-mannose-derived O-antigen building blocks biosynthesis | Super-Pathways |
| COLANSYN.PWY | -0.04415063059 | 0.01751043647 | 0.1756881841 | colanic acid building blocks biosynthesis | Super-Pathways |
| PANTOSYN.PWY | -0.04086930306 | 0.004019143793 | 0.09595114474 | pantothenate and coenzyme A biosynthesis I | Super-Pathways |
| PWY.5913 | -0.08105625381 | 0.03842318749 | 0.2322468221 | TCA cycle VI (obligate autotrophs) | TCA-VARIANTS |
| PWY.6123 | 0.01903853739 | 0.0249823896 | 0.1979187371 | inosine-5'-phosphate biosynthesis I | Nucleotide-Biosynthesis |
| PWY.7663 | 0.02001715026 | 0.03662202527 | 0.2290113368 | gondoate biosynthesis (anaerobic) | Lipid-Biosynthesis |

| | | | | | |
|---|---|---|---|---|---|
| PWY.5973 | 0.0227374266 | 0.02998662897 | 0.2073651631 | cis-vaccenate biosynthesis | Lipid-Biosynthesis |
| HISDEG.PWY | 0.1216791687 | 0.03668680945 | 0.2290113368 | L-histidine degradation I | Amino-Acid-Degradation |
| BIOTIN.BIOSYNTHESIS.PWY | 0.319047977 | 0.01976728007 | 0.1756881841 | biotin biosynthesis I | Super-Pathways |
| BIOTIN.BIOSYNTHESIS.PWY | 0.319047977 | 0.01976728007 | 0.1756881841 | biotin biosynthesis I | Cofactor-Biosynthesis |
| PWY.6519 | 0.3235450381 | 0.01984105858 | 0.1756881841 | 8-amino-7-oxononanoate biosynthesis I | Cofactor-Biosynthesis |
| PWY.5971 | 0.3269762344 | 0.01929039388 | 0.1756881841 | palmitate biosynthesis II (bacteria and plants) | Lipid-Biosynthesis |
| PWYG.321 | 0.3296549682 | 0.02014096333 | 0.1756881841 | mycolate biosynthesis | Lipid-Biosynthesis |
| PWY.7664 | 0.3301938468 | 0.02006855063 | 0.1756881841 | oleate biosynthesis IV (anaerobic) | Lipid-Biosynthesis |
| PWY0.862 | 0.3307684069 | 0.0202318048 | 0.1756881841 | (5Z)-dodec-5-enoate biosynthesis | Lipid-Biosynthesis |
| PWY.6282 | 0.3310216316 | 0.02035180229 | 0.1756881841 | palmitoleate biosynthesis I (from (5Z)-dodec-5-enoate) | Lipid-Biosynthesis |
| PWY.5989 | 0.3310964705 | 0.02052118987 | 0.1756881841 | stearate biosynthesis II (bacteria and plants) | Lipid-Biosynthesis |
| FASYN.INITIAL.PWY | 0.3314080798 | 0.02038061956 | 0.1756881841 | superpathway of fatty acid biosynthesis initiation (E. coli) | Super-Pathways |
| FASYN.INITIAL.PWY | 0.3314080798 | 0.02038061956 | 0.1756881841 | superpathway of fatty acid biosynthesis initiation (E. coli) | Lipid-Biosynthesis |
| P105.PWY | 0.4834897532 | 0.03732660268 | 0.2290113368 | TCA cycle IV (2-oxoglutarate decarboxylase) | TCA-VARIANTS |
